# Regulatory asymmetry in the negative single-input module network motif: Role of network size, growth rate and binding affinity

**DOI:** 10.1101/865527

**Authors:** Md Zulfikar Ali, Vinuselvi Parisutham, Sandeep Choubey, Robert C. Brewster

## Abstract

The single-input module (SIM) is a regulatory motif capable of coordinating gene expression across functionally related genes. We explore the relationship between regulation of the central autoregulated TF in a negatively regulated SIM and the target genes using a synthetic biology approach paired with stochastic simulations. Surprisingly, we find a fundamental asymmetry in the level of regulation experienced by the TF gene and its targets, even if they have identical regulatory DNA; the TF gene experiences stronger repression than its targets. This asymmetry is not predicted from deterministic modeling of the system but is revealed from corresponding stochastic simulations. The magnitude of asymmetry depends on factors such as the number of targets in the SIM, TF degradation rate (or growth rate) and TF binding affinity. Beyond implications for SIM motifs, the influence of network connectivity on regulatory levels highlights an interesting challenge for predictive models of gene regulation.

## Introduction

Gene regulatory networks are composed of an organisms’ genes connected based on the ability of some protein products (transcription factors or TFs) to alter the expression patterns of those genes. The networks, when viewed as a whole, are typically dense and interconnected and as such difficult to interpret (*1, 2*). The concept of network motifs, defined as overrepresented patterns of connections between genes and TFs in the network, helps to digest these large networks into smaller subgraphs with specific properties; each of these motifs can be interpreted as performing a particular “information processing” function that is determined by the connectivity and regulatory role of the genes in the motif (*2–6*).

The single-input module (SIM) is a network motif where a single TF regulates the expression of a set of genes, often including itself (Fig. **1A**). Typically, this group of genes have related functions and the purpose of this motif is to coordinate, in both time and magnitude, expression of these related genes (*6*). There are mounting examples, from diverse topics that range from metabolism (Fig. **1B**, (*7*)), stress response (Fig. **1C**, (*8, 9*)), development (*10–12*), and cancer (*13*), where temporal ordering of gene expression in the motif naturally follows the functional order of the genes in the physiological pathway. Mechanistically, it is thought that this ordering is set through differential affinity for the TF amongst the various target genes in the motif, where the strongest binding sites are the first ones to respond (and the last ones to stop responding) to a signal; this temporal patterning is referred to as “last-in first-out” (*6*). Due to the broad importance of these motifs, a quantitative understanding of how SIM modules can be encoded, designed and optimized, will be instrumental in gaining a deep and fundamental understanding of the spatial and temporal features of a diverse set of cellular phenomena.

**Fig. 1:**
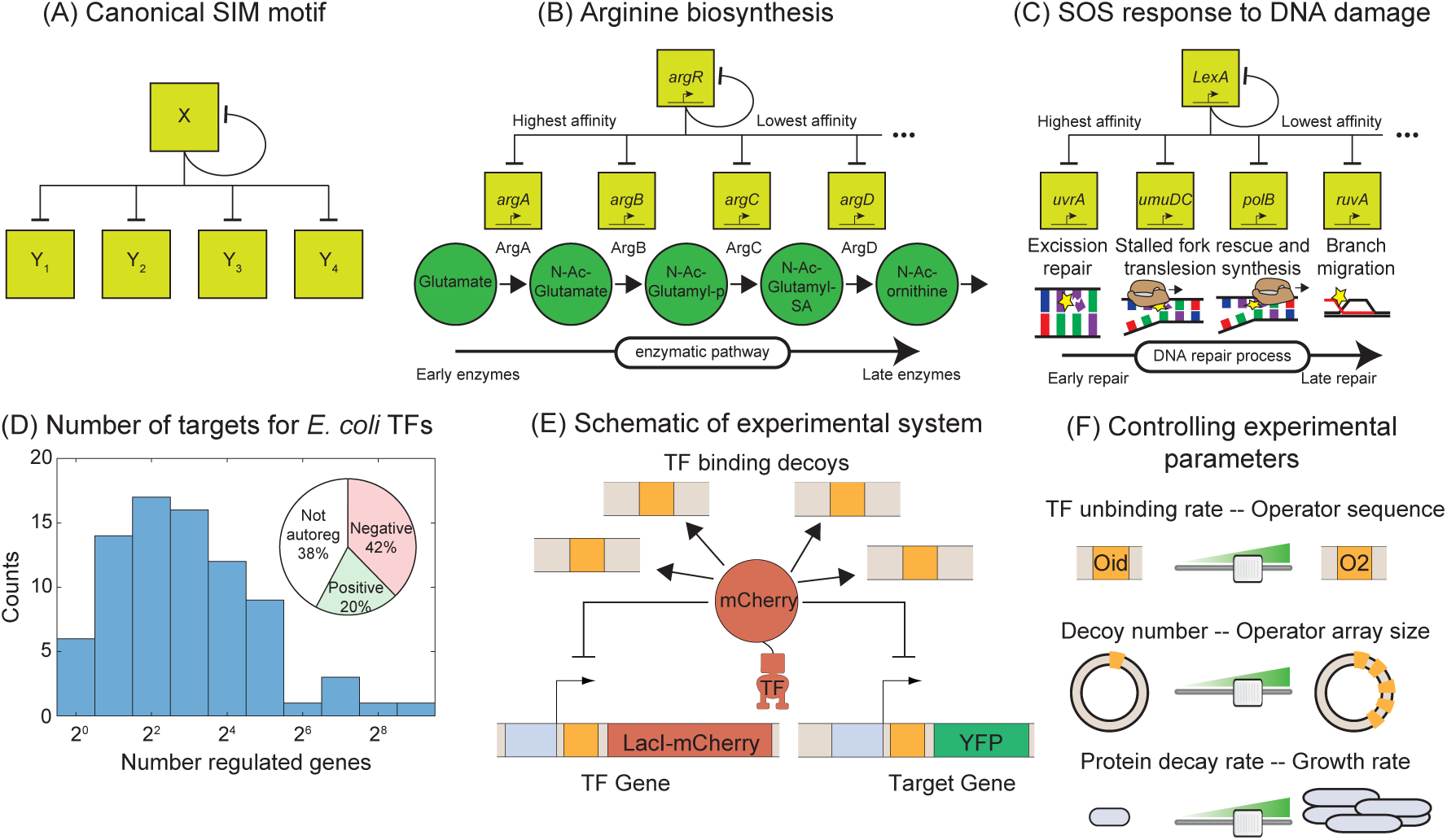
Synthetic approach to exploring the negative SIM motif. (**A**) Schematic of a canonical SIM motif: A single TF regulates itself and several other genes. (**B** and **C**) Examples of SIM motifs in *E. coli*. (**B**) ArgR is a transcriptional regulator of arginine biosynthesis. It auto-regulates itself and genes involved in different steps of arginine biosynthesis with precision in expression starting from the first enzyme of the pathway down to the last. This precise ordering is thought to originate from a corresponding ordering in TF binding affinities of the target genes. (**C**) LexA is the master regulator of SOS pathway and is actively degraded in response to DNA damage. LexA auto-represses itself and represses a set of other genes involved in DNA repair. In this case the early response genes have low affinity for the repressor while the late acting genes have high affinity, enabling temporal ordering of the response. (**D**) Histogram showing the number of known regulated genes for every TF in *E. coli*. Inset shows the percentage of TFs that are positive, negative or not autoregulated. (**E**) Schematic of the experimental model of a SIM motif used in this study. Here LacI-mCherry is the model TF and YFP is the protein product of the target gene. Decoys sites are used to control the network size by simulating the demand of other target genes in the SIM motif. (**F**) Representation of the tunable parameter space detailed in this study. We can systematically tune the TF unbinding rate, number of decoys and protein degradation rate in the experimental system and adjust these parameters accordingly in simulations.

To quantitatively explore the input-output relationship of the SIM motif, we use a synthetic biology approach that boils the motif down to its most basic components: an autoregulated TF gene, a sample target gene, and competing binding sites. Specifically, we use non-functional “decoy” binding sites to exert competition for the TF and mimic the demand of the other genes in the motif (which will depend on the size of the network, Fig. **1D** (*18*, *19*)). However, the demand for the TF could also stem from a litany of sources such as random non-functional sites in the genome (*14–17*) or non-DNA based obstruction or localization effects that transiently interfere with a TFs ability to bind DNA. Because of the design, our results do not depend on the nature of the TF competition. SIM TFs typically exert the same regulatory role on all targets of the motif (*18*). As such, in this work we will focus on a TF that is a negative regulator of its target genes and itself; this is the most common regulation strategy in *Escherichia coli* where roughly 60% of TF genes are autoregulated and almost 70% of those TFs negatively regulate their own expression (inset Fig. **1D**, (*18*)).

We used stochastic simulations of kinetic models (*19–22*), to predict how the overall level of gene expression depends on parameters characterizing cellular environment such as TF binding affinities and the number of competing binding sites. To test these predictions *in vivo*, we built a synthetic system with LacI as a model TF, and individually tune each of these parameters. Past work with LacI have demonstrated the ability to control with precision the regulatory function, binding affinity and TF copy number through basic sequence level manipulations (*23–30*); Here we use that detailed knowledge to inform our simulations which then guide our experiments (and vice versa).

Our approach reveals that the presence of competing TF binding sites can have counterintuitive effects on the mean expression levels of the TF and its target genes due to the opposing relationship between total TFs and free TFs (those not bound to a specific binding site). Furthermore, we find that the TF and target gene experience quantitatively different levels of regulation in the same cell, and with the same regulatory sequence. We show that this regulatory asymmetry is sensitive to features such as the degradation rate, TF binding affinity and the number of competing binding sites for the TF. Interestingly, regulatory asymmetry is not captured by a deterministic model of our stochastic simulation, which is based on mass action equilibrium kinetics and are widely used in predicting gene expression patterns and levels (including precise quantitative agreement for the promoter used in our study (*23, 25, 28, 29, 31*)). In fact, this deterministic model fails to accurately predict expression of either gene. However, the stochastic model makes accurate predictions that we confirm through *in vivo* measurements.

## Results

### Matching molecular biology with simulation methodology

We use a combination of theory and experimental *in vivo* measurements to study the interplay between TF, target, and additional binding sites of a negative autoregulatory SIM network motif. The basic regulatory system is outlined in Fig. **1E**. We use a kinetic model of the SIM motif to explore how the expression of the TF gene and one target gene depends on parameters such as TF binding affinity and number of other binding sites in the network (here modeled and controlled through competing, non-regulatory decoy sites (*32*)). In this model, the TF gene and target gene can be independently bound by a free TF to shut off gene expression until the TF unbinds. The two genes (TF-encoding and target) compete with decoy binding sites which can also bind free TFs. Each free TF can bind any open operator site with equal probability (set by the binding rate). The unbinding rate can be set individually for the TF gene, target gene and decoy sites and is related to the specific base pair identity of the bound operator site (*33–36*). We employ stochastic simulations to make specific predictions for how the expression level of the TF and target genes depend on the various parameters of the model. Furthermore, we translate these stochastic processes into a deterministic ODE model using equilibrium mass action kinetics (see SI section **S6**). This approach is prevalent in theoretical studies of gene expression because the equations can often be solved analytically and thus provide intuition for regulatory behavior of the system. Below we compare the stochastic and deterministic approaches to exploring the regulation of the SIM motif. A thorough discussion on how we chose the kinetic parameters of our model is presented in the methods section.

In experiments, the corresponding system is constructed with an integrated copy of both the TF (LacI-mCherry) and target gene (YFP) with expression of both genes controlled by identical promoters with a single LacI binding site centered at +11 relative to their transcription start sites (*23, 28*). As demonstrated in Fig. **1F**, decoy binding sites are added by introducing a plasmid with an array of TF binding sites (between 0 to 5 sites per plasmid) enabling control of up to roughly 300 binding sites per cell (for average plasmid copy number measured by qPCR, see methods and SI Fig. **S3**). TF unbinding rate is controlled by changing the sequence identity of the operator sites; the binding sequence assessed in this study include (in order of increasing affinity) O2, O1 and Oid. The decoy binding site arrays are constructed using the Oid operator site. We quantify regulation through measurements of fold-change (FC) in expression which is defined as the expression level of a gene in a given condition (typically a specific number of decoy binding sites) divided by the expression of that gene when it is unregulated. For the target gene we can always measure unregulated expression simply by measuring expression in a LacI knockout strain. However, it is challenging to measure unregulated expression for the autoregulated gene. For autoregulation this unregulated expression can be measured by exchanging the TF binding site with a mutated non-binding version of the site. For O1 there is a mutated sequence (NoO1v1 (*30*)) that we have shown relieves repression of the target gene comparable to a strain expressing no TF (see SI Fig. **S4B**) which allows us to calculate fold-change even for the autorepressed gene. Despite testing many different mutated sites and strategies, we could not find a corresponding sequence for O2 and Oid so we focus primarily on studying a TF gene regulated by O1 (see SI text **S4** for more discussion).

### Decoy sites increase expression of the auto-repressed gene and its targets

We first investigate the negatively regulated SIM motif where the TF and target gene have identical promoters and TF binding sites (O1) and the number of (identical) competing binding sites are varied systematically (schematically shown in Fig. **1E, F**). Simulation and experimental data for fold-change of the TF gene as a function of number of decoys is shown in Fig. **2A** as red lines (simulation) and red points (experiments). We find that increasing the number of decoy sites increases the expression of the auto-repressed TF gene monotonically. To interpret why the TF level increases, in Fig. **2B** we plot the number of “free” TFs in our simulation (defined as TFs not bound to an operator site) as a function of decoy site number. The solid line demonstrates that on average, despite the increased average number of TFs in the cell, the number of unbound TFs decreases as the number of competing binding sites increases. Therefore, because the number of available repressors decreases, the overall level of repression also decreases and thus the mean expression of the TF gene rises.

**Fig. 2:**
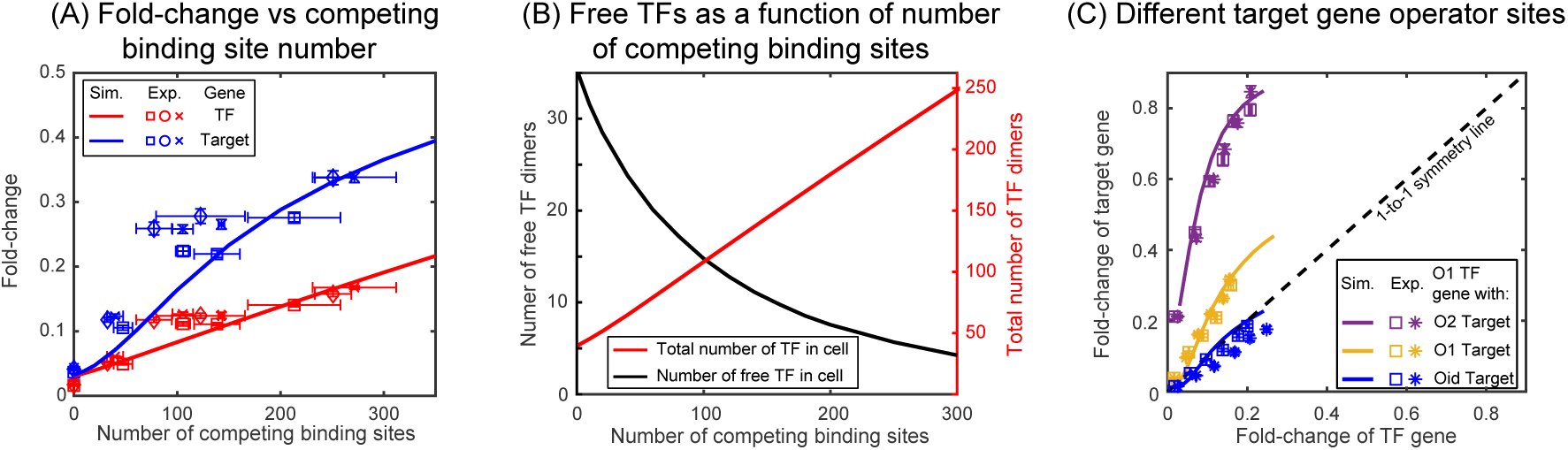
Fold-change in target and TF genes with network size. (**A**) Fold-change in the expression level of both the autoregulated gene (red) and the TF’s target gene (blue) as a function of the number of competing binding sites present. (**B**) Increasing the number of competing binding sites increases the expression of both the TF (red line) and target genes by lowering the overall number of free TFs (black line). (**C**) Fold-change in the target gene versus fold-change in the TF gene. Each data point is a population average in TF and target expression across hundreds of cells with a given number of competing binding sites. In all cases the TF gene is regulated by an O1 binding sites whereas the target is regulated by (in order of weakest binding to strongest binding): O2 (purple), O1 (yellow) or Oid (blue). Simulation data is shown as solid curves whereas open squares, circle and stars are experimental data. Error bars in experimental data are standard error of the mean obtained using bootstrapping method.

Now we consider the regulation of a SIM target gene which is regulated by an O1 binding site. In this case, the target promoter and TF promoter are identical. In Fig. **2A**, the expression of the target gene is shown as blue points (experiments) and blue lines (simulation) for the SIM motif with different numbers of decoy TF binding sites. Just as in the case of the TF gene, we once again see that the expression of the target gene increases as more decoy binding sites are added even though the total number of TFs is also increasing (red points and line). Qualitatively, we expected this result since the free TF number is expected to decrease (Fig. **2B**) and, in turn, the expression of any gene targeted by the autoregulated repressing TF will increase. While the mechanism is more obvious in this controlled system, it is important to note that this is a case where more repressors correlate with more expression of the repressed gene. It is easy to see how this relationship could be misinterpreted as activation in more complex *in vivo* system if the competition level of the TF is (advertently or otherwise) altered in experiments.

### Asymmetry in gene regulation between TF and target genes

The stochastic simulations and experimental data in Fig. 2**A** reveal an intriguing detail: Even when the regulatory region of the auto-repressed gene and the target gene are identical, we find that the expression (fold-change or FC) is higher for the target gene, raising the question of how two genes with identical promoters and regulatory binding sites in the same cell can have different regulation levels. Interestingly, this finding stands in sharp contrast to what the corresponding deterministic modeling predicts, the expressions of the target and TF genes are expected to be identical when the corresponding regulatory regions are the same (see SI Fig. **S7C**, (*37*)). The asymmetry in regulation, as predicted by the stochastic simulations, is shown explicitly in Fig. **2C**, where we plot the expression of the target gene against the expression of the TF gene. In this figure, the data points are derived from measurements made in six different competition levels (from 0 to 5 decoy binding sites per plasmid). Each data point represents the average expression level of each gene for a given number of competing binding sites. The lines represent the same quantity calculated by simulation. The intuitive expectation, further bolstered by the deterministic model predictions, that identical promoters (yellow data, Fig. **2C**) should experience identical levels of regulation would suggest that the data fall on the black dashed one-to-one line. However, for both simulations and experiments of this system the TF gene is clearly more strongly regulated than the target gene subject to identical regulatory sequences.

To examine the extent of asymmetry in this system, we adjust the target binding site to be of higher affinity (Oid, blue lines and data points in Fig. **2C**) or weaker (O2, purple lines and data points in Fig. **2C**). Clearly, this should change the symmetry of the regulation, after all the TF binding sites on the promoters are now different and symmetry is no longer to be expected. The experiments and simulations once again agree well. However, when Oid regulates the target gene and O1 regulates the TF gene, the regulation is now roughly symmetric despite the target gene having a much stronger binding site; in this case, the size of the inherent regulatory asymmetry effect is on par with altering the binding site to a stronger operator resulting in symmetric overall regulation of the genes.

### Mechanism of asymmetric gene regulation

The difference in expression between the TF and its target can be understood by studying the TF-operator occupancy for each gene, drawn schematically in Fig. **3A**. This cartoon shows the four possible promoter occupancy states of the system: (1) both genes unbound by TF, (2) target gene bound by TF, TF gene unbound, (3) TF gene bound by TF, target gene unbound, and (4) both genes bound by TF. It should be clear that state 1 and state 4 cannot be the cause of asymmetry; both genes are either fully on (state 1) or fully off (state 4). As such the asymmetry must originate from differences in states 2 and 3. In state 2, the TF gene is “on” while the target gene is fully repressed and in state 3 the opposite is true. Since we know that the asymmetry appears as more regulation of the TF gene than the target gene, then it must be the case that the system spends less time in state 2 than in state 3. There are two paths to exit either of these states: unbinding of the TF from the bound operator or binding of the TF to the free operator. Since unbinding rate of a TF is identical for both promoters in our model, the asymmetry must originate from differences in binding of free TF in state 2 and in state 3; specifically state 2 must have an (on average) higher concentration of TF than state 3. This makes sense since the system is still making TF in state 2, while production of TF is shut off in state 3. Fig. **3B** validates this interpretation as we can see that state 2 has on average more free TFs than state 3, and as a result, the system spends less time in state 2 than in state 3 in our simulations. As such, the asymmetry comes from the fact that the two genes, despite being in the same cell and experiencing the same average intracellular TF concentrations, are exposed to systematically different concentrations of TF when the TF and target gene are in their respective “active” states. To quantify regulatory asymmetry, we define asymmetry as the difference in fold-change of the target and TF gene (asymmetry =FC_target_-FC_TF_). In our simulations we find that asymmetry is exactly equal to the difference in time spent in state 3 and state 2, for any condition or parameter choice (Fig. **3C**).

**Fig. 3:**
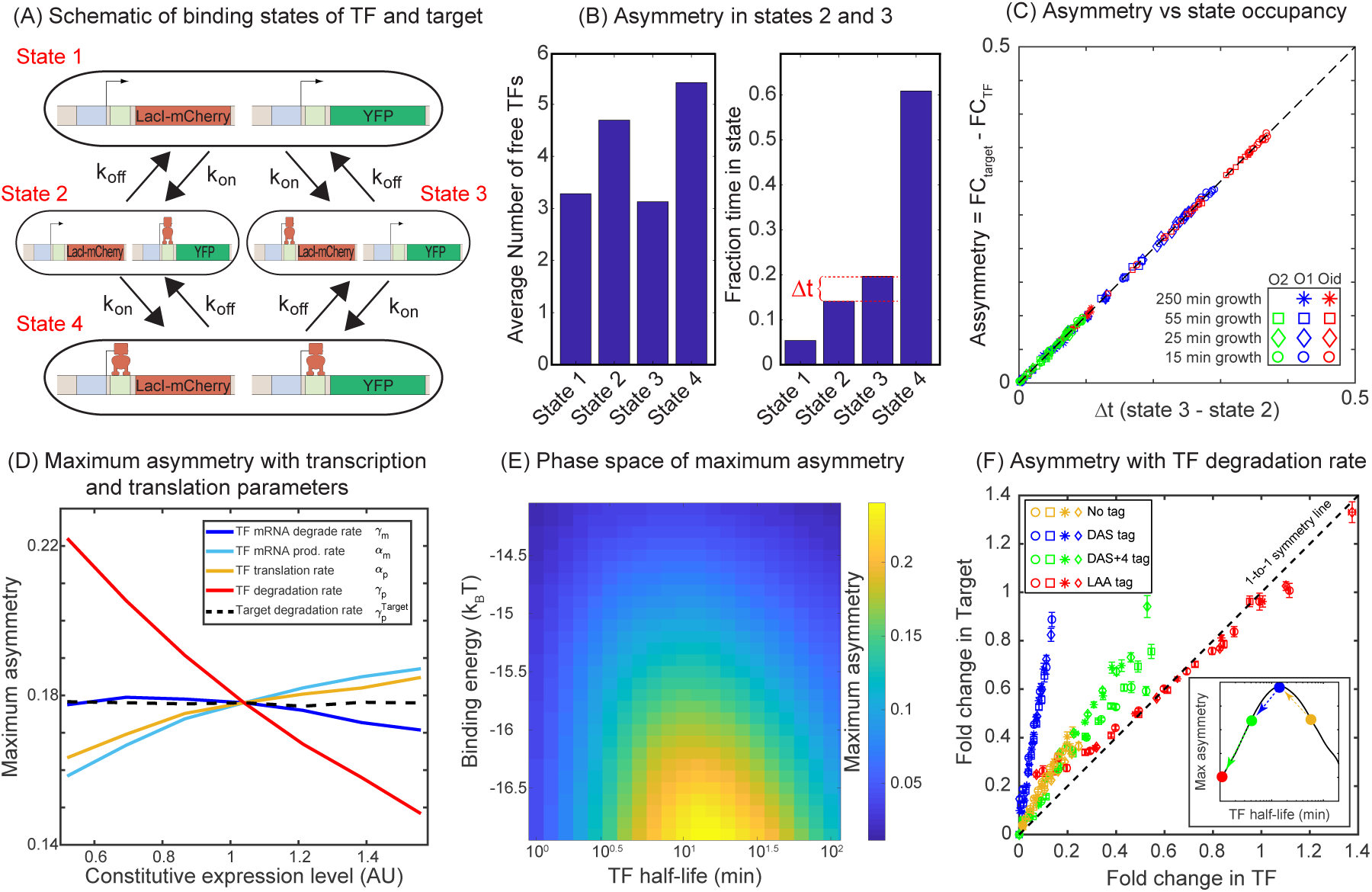
Mechanism of regulatory asymmetry. (**A**) Schematic of the TF-operator occupancy with their corresponding transition rates. The *k*_on_ for transition from state 1 to state 2 or state 3 will be identical and hence cannot account for the asymmetry. State 2 and state 3 on the other hand, will encounter a difference in the free TF concentration and hence the *k*_on_ for transition from one of these states to state 4 will be different; thus, accounting for the asymmetry in expression between the TF and the target. (**B**) Plot showing the average number of free TFs in different states and fraction of time cells spends in each of the given state in the simulation. (**C**) Plot showing asymmetry as a function of fractional time difference between state 2 and state 3. (**D**) Exploring the model parameters of the TF (mRNA production and degradation; protein production and degradation) that could influence the asymmetry between the TF and the target. Tuning the protein degradation rate (red line) has the maximum influence on the asymmetry between the TF and its target gene. (**E**) Heat map showing the phase space of maximum asymmetry as a function of binding affinity for the TF and its half-life. (**F**) Tuning the TF degradation rate influences the extent of asymmetry observed in the SIM module. Yellow points correspond to the system with no degradation tags; Blue points correspond to degradation by a “weak” or “slow” tag (DAS tag); Green points correspond to a slightly faster tag (DAS+4); Red points corresponds to a very fast tag (LAA tag). Inset shows schematic of the expected maximum asymmetry as degradation rate of the TF is increased.

According to the above proposed mechanism, the regulatory asymmetry stems from differences in the cellular TF concentration when the TF is bound to the target versus when it is bound to the autoregulatory gene, as such we expect that binding affinity will play a central role in setting asymmetry levels. However, there are many parameters associated with the production and decay of TF and target mRNA and protein which could also influence the asymmetry. To reveal which (if any) of these parameters is important to asymmetry, Fig. **3D** shows our theory predictions for the maximum asymmetry (the maximum value of asymmetry found as competing site number is controlled, see SI Fig. **S8**) as these production and degradation parameters are tuned. First, we find that tuning the rates of target gene production and decay has almost no effect on asymmetry (black dashed line for target protein degradation rate, others not shown). On the other hand, for TF production and decay each parameter has some effect on asymmetry. However, we find that the biggest driver of asymmetry in this set of parameters is the protein degradation rate (red line). As such, we focus on two crucial parameters that control the asymmetry: TF binding affinity and TF degradation rate. In Fig. **3E** we show a heat map of the maximum asymmetry as a function of the rate of protein degradation and binding affinity of the TF. We see from this figure that strong binding produces enhanced asymmetry, and the degradation rate displays an interesting intermediate maximum in asymmetry – degradation that is too fast, or too slow will not show asymmetry. A maximum asymmetry is expected for TF lifetimes between 10 and 100 minutes. Crucially, this maximum coincides with typical doubling time of *E. coli* (which sets the TF half-life (*38, 39*)) and thus asymmetry is most relevant in common physiological conditions

### Dependence of regulatory asymmetry on TF degradation and binding affinity

To experimentally test the theory predictions for the role of TF degradation in setting regulatory asymmetry, we introduced several ssrA degradation tags to the LacI-mCherry in our experiments (*40*). The data, shown in Fig. **3F** includes degradation by a “weak” or “slow” tag (DAS with a rate of 0.00063 per minute per enzyme (*41*), blue points), a slightly faster tag (DAS+4 with a rate of 0.0011 per minute per enzyme (*41*), green points) and a very fast tag (LAA tag with a rate of 0.21 per minute per enzyme (*41*), red points). In addition, the data without a tag is shown as yellow points. Here we see that the slowest tag (blue points) introduces strong asymmetry. However, for the next fastest tag (green points) we see a significant decrease in asymmetry and the level of regulatory asymmetry is similar to what is seen in the absence of tags (yellow points). Finally, the fastest tag (red points) shows no asymmetry at all. It is worth pointing out that the qualitative order of degradation rates in these experiments can be inferred from how far the data “reaches,” faster degradation will lead to higher overall fold-changes for a given competition level. Importantly, controlling the protein degradation rate through this synthetic tool agrees with our model predictions, although the actual *in vivo* protein degradation rates are difficult to estimate from tag sequence alone, the asymmetry follows the expected trends based on the known (and observed) effectiveness of each tag (see schematic inset Fig. **3F**).

In the absence of targeted degradation, the degradation rate of most protein in *E. coli*, is naturally set by the growth rate. According to the model predictions in Fig. **3E**, the asymmetry should be highest for fast growing cells (roughly 20-minute division rates) and decrease (or vanish) for very slow growing cells. To test this, we take the system with O1 regulatory binding sites on both the target and the TF promoter (yellow data in Fig. **2C** grown in M9 + glucose, 55-minute doubling time) and grow in a range of doubling times between 22 minutes (rich defined media) up to 215 minutes (M9 + acetate) (see SI Fig**. S2A**). Importantly, when we change the growth rate, other rates such as the transcription and translation rates will also be impacted (*42, 43*), while these parameters will change features of the asymmetry curve (see Fig. **3D**), the qualitative ordering and features of the asymmetry are not expected to be impacted (see SI Fig. **S6**). The data for these growth conditions is shown in Fig. **4A**. As predicted, faster growing cells show more regulatory asymmetry and slower growing cells show little-to-no regulatory asymmetry. We also test the role of growth rate in asymmetric regulation when O2 (a lower affinity site) and Oid (a higher affinity site) are used as the regulatory binding sites instead of O1. This data is shown in Fig. **4B** (O2) and **4C** (Oid). As discussed above, we could not find a suitable mutant for O2 and Oid that both relieved regulation from LacI and completely restored the expression of target gene (see SI text **S4**.). This means we cannot explicitly measure the 1-1 correlation between the two axes in our data when using O2 or Oid for the TF gene. To this end, we find this correspondence by fitting the glucose data to our simulation of the same system and use that value to normalize all other growth rates for that operator. Despite this complication, it is clear that O2 regulation is symmetric at all studied growth rates while Oid regulation is asymmetric for all growth rates with faster growth rates appearing more asymmetric.

**Fig. 4:**
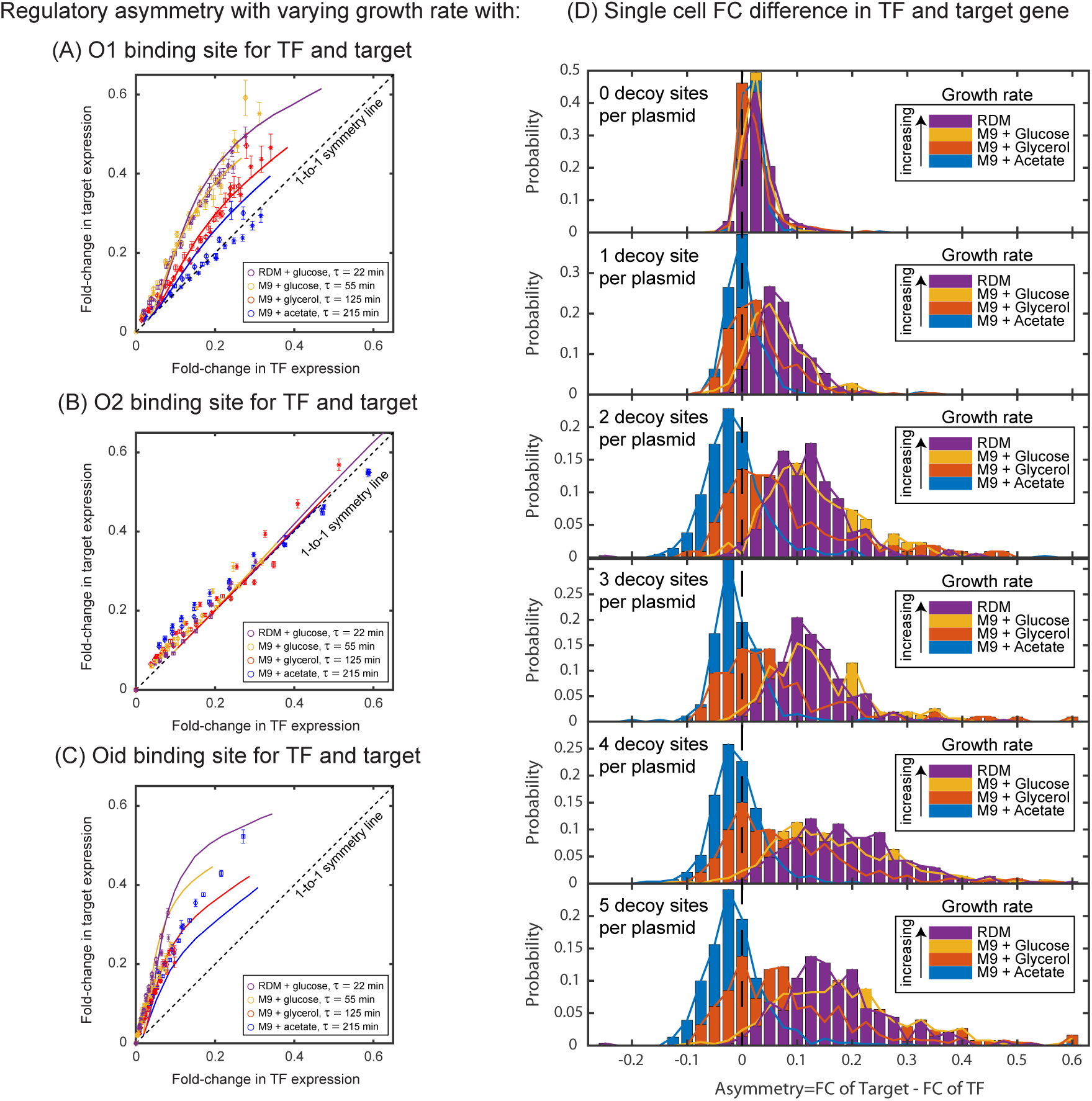
Dependence of regulatory asymmetry on growth rate. Measurement of asymmetry in different media as a function of TF binding energy: O1 (**A**), O2 (**B**), Oid (**C**). The division time (τ) is varied between 22 minutes up to 215 minutes. (**A**) For O1, the asymmetry decreases with slower division rates and agrees well with the simulation predictions. (**B**) For the weak O2 site, no asymmetry is seen at any growth rate. (**C**) For the strongest site Oid asymmetry is present at every growth rate although the magnitude of asymmetry still orders roughly by growth rate. (**D**) Histograms of single-cell asymmetry in expression of the TF and target gene regulated by O1 binding site in these 4 growth rates. Panels from top to bottom represent increasing the level of competition for the TF.

Importantly, the regulatory asymmetry is not due to a small population of outliers, bimodality or any other “rare” phenotype. In Fig. **4D**, we show a histogram of single cell asymmetry values (defined as asymmetry = FC_Target_– FC_TF_) for each condition. As can be seen, expression in each media condition are roughly symmetric for most cells at the lowest competition levels (top panel). However, as competition levels are increased, the fast-growing conditions shift to higher asymmetry levels; strikingly at the highest growth rate almost every single cell is expressing target at a higher level than TF (bottom panel).

### Divergence from deterministic model solutions

In this study, both the TF gene and target gene have identical regulatory regions; both genes are regulated by a single repressor binding site immediately downstream of the promoter. This regulatory scheme is often referred to as “simple repression” (*28, 44, 45*). For regulation of this kind, we can derive the expected fold-change using the deterministic modeling approach described here (see SI text **S6**). Under this framework, we find that regardless of the network architecture (autoregulation, constitutive TF production, number of competing sites, *etc.*), the fold-change is expected to follow a simple scaling relation,

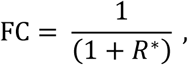

where,

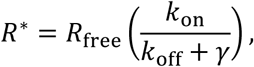

where *R*_free_ is the number of free (unbound) TFs and *k*_on_/(*k*_off_ + *γ*) is the affinity of the specific TF binding site. Although the free TF concentration is inherently difficult to measure experimentally, it has previously been shown that in the thermodynamic framework *R*_free_ is a calculable quantity (where it is directly related to the TF “fugacity”). The fugacity is determined from details that will alter TF availability such as total number of TFs, number of decoy binding sites, TF binding affinities and inducer concentrations (*23, 44, 46–49*). The advantage of this approach is that for experimental data the effective concentration is calculable from basic measurable parameters of the system. Importantly the two approaches (using fugacity or free TF) yield identical results.

Fig. **5A** shows a collection of experimental measurements (adapted from (*50*)) where the free concentration of TF is varied through any of these parameters (binding affinities, total TF number, inducer concentration, *etc.*) and the resulting fold-change is measured as a function of the free TF concentration; the predictions of the thermodynamic model are extremely robust to these perturbations and the collapse of this data demonstrates that the theory has identified the “natural variable” (free TF concentration) of the system in agreement with the deterministic expectation (black dashed line Fig. **5A**). In the studies comprising the data of Fig. **5A** and in other similar quantitative studies, the TF is expressed either constitutively or from an inducible promoter controlled by a second repressor, (*23, 25, 28, 51*). However, in the case of negative autoregulation and the resulting regulatory asymmetry, the above fold-change relationship obviously cannot hold for both the TF and target gene, but it is unclear where the departure from this relationship occurs. In fact, it has previously been shown that the binding probability of a TF to an autoregulatory gene can deviate from the deterministic solution (*52–54*), so it is reasonable to expect that this is the root of the asymmetry. In Fig. **5B**, we show simulation data for the fold-change versus number of scaled-free TFs (*R\**) for the autoregulatory gene (red line) and its target gene (blue line) with O1 (Fig. **5B**), O2 (Fig. **5C**) and Oid (Fig. **5D**) binding sites, where we are changing the number of free TFs by tuning the number of competing binding sites. In each plot, we also show simulations for the fold-change of a single target gene with a TF undergoing constitutive (constant in time) expression where the TF is controlled by either changing the expression level of the TF (purple stars) or adding competing binding sites while maintaining a set constitutive expression level (purple circles). In both cases, where TFs are made constitutively, the simulation data agrees well with the deterministic model predictions (although the strongest binding site, Oid, begins to show some divergence). However, for the autoregulatory circuits, we find that for strong binding sites (O1 and Oid) neither the target nor the TF gene follow the deterministic solution (black dashed line), and surprisingly, the target gene deviates even more strongly from the deterministic solution. Thus, the asymmetry is not a result of one gene diverging from the deterministic solution but rather both genes diverging in different ways. As expected, when the binding is weak (O2), both the target and TF gene converge to the deterministic solution.

**Fig. 5:**
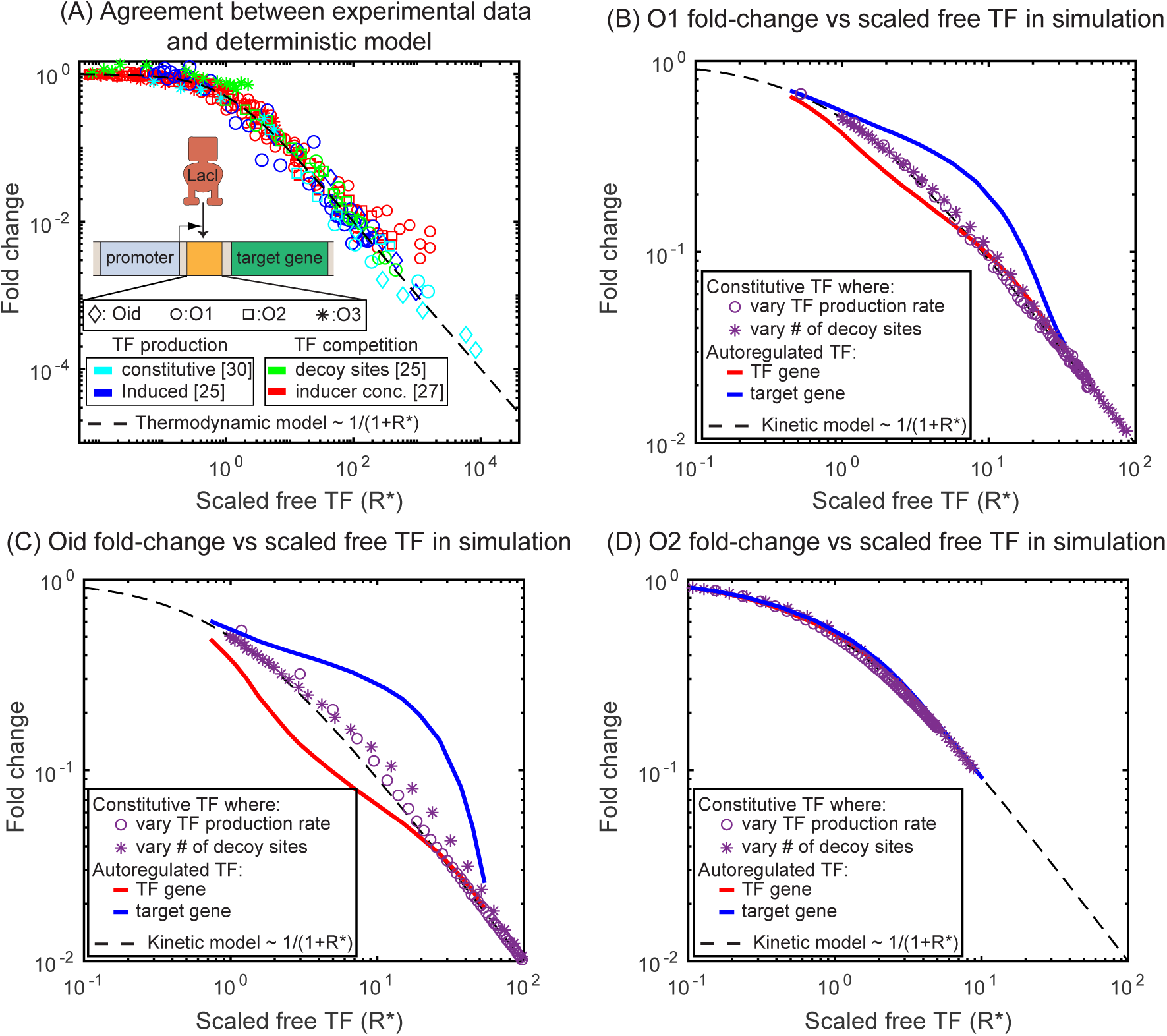
Comparison of SIM motif fold-change data to deterministic model predictions. (**A**) Fold-change vs scaled free TF in the thermodynamic model for a collection of simple repression data where free TF is controlled through a diverse range of mechanisms. The data collapses to the deterministic model predictions. (**B**-**D**) Fold-change vs scaled free TF in simulations using the actual free TF measured in simulation. The data for a constitutive expressed TF where free TF is varied by changing TF production rate (purple circles) or number of decoy sites (purple stars) collapses to the deterministic solution, however, the regulation of genes in the SIM motif (target: blue line, TF gene: red line) both diverge from the deterministic solution in opposing ways, giving rise not only to asymmetry but a disagreement with deterministic modeling for both genes.

## Discussion

The single input module (SIM) is a prevalent regulation strategy in both bacteria (*18, 55*) and higher organisms (*56–58*). While the role of TF autoregulation (positive and negative) has been extensively studied (*59–66*), the focus here is on the combined influence of an autoregulated TF and its target genes and how the shared need for the TF influences the quantitative features of its regulatory behaviors. We find that there is a fundamental asymmetry in gene regulation that can occur in the SIM regulatory motif. This asymmetry is not related to distinctions in the biological processes or an unexpected difference in our *in vivo* experiment, but rather an inherent asymmetry originating from the way the motif itself is wired. Although two identical promoters are in the same cell with the same average protein concentrations, they experience distinct regulatory environments. This is particularly relevant for the SIM motif because the primary function of the motif, organizing and coordinating gene expression patterns, operates on the premise of differential affinities amongst target genes; here we have shown that the TF gene has an inherent “affinity advantage” due to being exposed to systematically higher TF concentrations than its target genes.

Regulatory asymmetry is intrinsic to the negative SIM motif even in the absence of decoys, but it can be greatly exacerbated by competing TF binding sites. Due to the promiscuous nature of TF binding, this highlights the importance of considering not just the “closed” system of a TF and a given target but also the impact of other binding sites (or inactivating interactions) for the TF in predicting regulation as well as the regulatory motif at play in the system. In our system, the magnitude of the asymmetry is enough to compensate for swapping the wild-type proximal O1 LacI binding site on the target gene with the “ideal” operator Oid.

The cause of this asymmetry is a systematic difference in the TF concentration when the TF gene is active compared to when the target gene is active. As such, asymmetry is magnified by anything that enhances this concentration difference. Here we have identified TF binding affinity and TF degradation rate (controlled both directly and through modulating growth rate) as primary drivers of asymmetry in this motif. Although the relationship between growth rate and expression levels is well established (*42, 43, 67–69*), effects such as this add a layer of complexity to this relationship.

In studies of quantitative gene regulation, the typical goal is to predict the output of a gene based on the regulatory composition of that gene’s promoter and the number and identity of regulatory proteins. This work clearly presents a challenge for the drive to “read” and predict regulation levels from the promoter DNA alone, in this case the regulatory motif is responsible for altering the observed regulation and must be considered as well. It has previously been demonstrated that features of a transcript can impact its regulation by effects such as targeted degradation, stabilization or posttranslational modification and regulation (*70*), it is important to point out this is a distinct phenomenon that does not operate through an enzymatic process but rather is a fundamental feature of the network.

Finally, here we demonstrate regulatory asymmetry using a specific (but common) regulatory motif. The broader point that specific genes can be exposed to systematically different levels of regulatory TFs even in the absence of specific cellular mechanisms such as cytoplasmic compartmentalization, protein localization or DNA accessibility is likely more broadly relevant. Understanding and quantifying these mechanisms can be an important piece towards improving our ability to predict and design gene regulatory circuits.

## Acknowledgments

We wish to thank Rob Phillips, Griffin Chure, Manuel Razo-Mejia, Amir Mitchell, Job Dekker, Marian Walhout, and Michael Lee for helpful discussions. We thank Dr. Jeffrey Bailey for providing us with qubit for DNA quantification. We thank Kenan Murphy for his valuable suggestions on protein degradation tags.

## Author contributions

MA did the computational analysis; VP performed all the experiments; SC provided the necessary supervision for the computational setup; RB conceptualized the experiments and drafted the manuscript. All data and codes are backed up in UMASS server and are readily available on request. We declare no conflict of interest.

