## Supplemental materials for "Regulatory asymmetry in the negative single-input module network motif: Role of network size, growth rate and binding affinity"

### Supplementary Materials for Inherent regulatory asymmetry in the negative SIM motif: Dependence on network size, growth rate and binding affinities

Md Zulfikar Ali[a,b,1], Vinuselvi Parisutham[a,b,1], Sandeep Choubey[c], Robert C. Brewster[a,b,*]

[a]*Program in Systems Biology, University of Massachusetts Medical School, 368 Plantation St., Worcester, MA 01605*
[b]*Department of Microbiology and Physiological Systems, University of Massachusetts Medical School, 368 Plantation St., Worcester, MA 01605.*
[c]*Max Planck institute for the Physics of Complex Systems, Nothnitzer strasse 38, 01187 Dresden, Germany*

---

**This PDF file includes:**

Materials and Methods

Supplementary Text

Figs. S1 to S9

Tables S1 to S3

[1]Authors contributed equally to this work.

#### Materials and Methods

##### Bacterial Strains

All strains used in this study are constructed from the parent strain *E. coli* HG105 which is MG1655 with the *lac* operon deleted (MG1655 Δ*lacIZYA*). Auto-regulated TF (*lacI-mCherry*) is expressed from the *ybcN* locus and the TF-repressed target (*yfp*) is expressed from the *galK* locus with identical promoter sequence for both the TF and the target. Decoys are introduced on the pZE plasmid. In order to tune the degradation rate of the TF, three different ssrA tags were added to the C-terminus of the LacI-mCherry fusion protein. The tags used in this study are wildtype LAA tag (AANDENYALAA), DAS tag (AANDENYADAS) and DAS+4 tag (AANDENYSENYADAS) *(40)*. For protein degradation tag experiments with LacI-mCherry fusion protein HG105 with $\Delta sspB$ knockout is used as a parent strain. Primers used in this study are listed in Table **S3**.

##### Microscopy

Bacterial cultures are grown overnight in 1 mL of LB in a 37°C incubator shaking at 250 rpm. Unless otherwise stated cultures grown overnight are diluted $2.5 \times 10^3$ fold into 1 mL of fresh M9 minimal media supplemented with $0.5\%$ of one of the three different carbon sources (Glucose, Glycerol or Acetate), allowed to grow at 37°C until they reach an OD600 of 0.2 to 0.4 (0.1 for acetate) and harvested for microscopy. Cells are diluted 1:3 in 1X PBS (in order to obtain isolated cells in microscope images) and 1 $\mu$L is spotted on a $2\%$ low melting agarose pad (Invitrogen $\#16520050$) made with 1X PBS. Rich defined media (RDM, Teknova $\#M2105$) is prepared according to the manufacturers instruction. Cells grown in RDM are cross-linked with para-formaldehyde before imaging to prevent shrinkage and osmotic shock to the cells. An automated fluorescent microscope (Nikon TI-E) with a heating chamber set at $37^\circ$C is used to record multiple fields per sample (between 8-12 unique fields of view) resulting in roughly 500 to 1000 individual cells per sample.

##### qPCR measurements for average plasmid copy number

We performed qPCR measurements in order to quantify the average copy number of the pZE plasmid. Cells are grown as described for microscopic analysis and diluted 1:200 in Qiagen P1 lysis buffer and allowed to sit on ice. Meanwhile, cells are plated at $10^{-5}$ dilutions on fresh LB plates in order to determine the colony forming units per mL (CFU/mL). 25 $\mu$L of the lysate is diluted with 25 $\mu$L of 1X PBS and allowed to sit for 5 minutes. The cells are then diluted 1:100 into 1X cut smart buffer from NEB. 20 $\mu$L of the mixture is incubated with $0.5$ $\mu$L of HindIII restriction enzyme for 30 minutes at 37°C followed by heat inactivation at $80^\circ$C for 20 minutes. The mixture is further diluted 1:10 and 4.2 $\mu$L is used as a template in a 20 $\mu$L qPCR reaction mixture. The pZE-1XOid plasmid is purified using the Qiagen Plasmid Medi Prep kit and quantified using the Qubit dsDNA assay kit. A standard curve is then prepared by diluting pZE-1XOid plasmid from $10^8$ copies down to 10 copies. The average copy number of the decoy plasmid per cell is computed by comparing the cT of the sample to the standard curve and dividing by the number of cells in the sample.

#### Simulation methodology

To model the experiments and study the effect of decoy sites on the expression of a target gene regulated by a negatively autoregulated TF gene, we develop a simple model of the experimental system. In our model the auto-regulatory gene produces a protein ($X$) which forms a TF dimer ($Z$). We explicitly modeled TF as a dimer to incorporate the fact that LacI acts as a dimer in our experimental system (the LacI-mCherry construct lacks the tetramerization domain (*71*)). Dimerization and de-dimerization steps occur at the rate $k_p$ and $k_m$, respectively. The TF binds to its own promoter ($P_{\text{TF}}$), to the promoter of the target gene ($P_{\text{target}}$), and to the decoy sites ($N$) with a constant rate $k_{\text{on}}$ per free-TF per unit time. The off rate of the bound TF ($k_{\text{off}}$, the unbinding rate) depends on the sequence identity and can be different for different promoters. A bound TF unbinds from the promoters of the TF and target, and from the decoy sites at a rate $k_{\text{off,TF}}$, $k_{\text{off,target}}$, and $k_{\text{off,decoy}}$ per unit time, respectively. A TF-free promoter produces an mRNA at the rate $\beta$ which is then translated into a protein at a rate $\alpha$. The mRNA and the proteins are degraded at the rate $\gamma_m$ and $\gamma$, respectively. We assume that all proteins (free protein, TF bound to promoter and TF bound to decoy sites) degrades with the same rate. Typically, the proteins in *E. coli* are very stable with protein half-life greater than the cell cycle and the dominant contribution to degradation comes from the dilution due to cell division. The degradation rate is thus given by $\gamma = ln(2)/\tau_1 + ln(2)/\tau_2$, where $\tau_1$ and $\tau_2$ are protein half-life and cell division time, respectively. The set of reactions describing the model above are listed in Table **S1A**.

We implement the simulations for stochastic reaction systems using Gillespies algorithm (*19*) and in C programming. Each simulation is run for sufficiently long time ($\sim 10^6$ s) to reach a steady state. Typically, for the rates used in this paper the steady state is achieved in $10^5$ s or less (see SI Fig. **S7A** for a sample time trace). Data for steady state distributions (TF and target protein) are then recorded by sampling over time with a time interval ($T_{\text{S}}$) long enough for the slowest reaction to occur 20 times on average ($T_{\text{S}} = 20$ over rate for slowest reaction). Mean protein numbers in steady state for fold-change are calculated using at least $10^5$ data points for each single run.

#### Kinetic parameter estimation

To compare the results from experiments with our simulations we are required to find values for the kinetic on and off rate of LacI for different operator sites (Oid, O1 and O2), the transcription and translation rates, mRNA degrdation rate, and the growth rates in different media. We directly measure growth rate for different media in our experiment (see SI text **S2**). The on and off rates are related to the binding energy ($\Delta\epsilon$) through,

$$\frac{k_{\text{on}}}{k_{\text{off}} + \gamma} = \frac{e^{-\Delta\epsilon/k_{\text{B}}T}}{N_{\text{ns}}} \tag{S1}$$

where $N_{\text{ns}} \sim 5 \times 10^6$ bps is the number of non-specific binding sites in the genome (which we take as the total number of bases) (*45*), $k_{\text{on}}$ is the binding rate per free TF per unit time, $k_{\text{off}}$ is the unbinding rate per unit time and $\gamma$ is the decay rate of the TF. Experimental measurements of $\Delta\epsilon$ have been reported in many repeated experiments (*23,25,28*) and thus we constrain our choice of $k_{\text{on}}$ and $k_{\text{off}}$ such that we obtain affinities consistent with these measurements. Taking one data set (O1 regulated TF and O1 regulated target grown in glucose), we use maximum likelihood analysis to obtain the rates by varying $k_{\mathrm{on}}$ in a range 0.0015-0.003 $\mathrm{s}^{-1}$ (which sets the corresponding value of $k_{\mathrm{off}}$ to give $\epsilon_{\mathrm{O1}} = 15.3\ k_{\mathrm{B}}T$ ), $\gamma_m^{-1}$ in a range of $30 - 90$ s, $\beta$ in a range of 0.1-0.3 $\mathrm{s}^{-1}$, and choosing $\alpha$ such that the constitutive number for the TF protein is in the range of $1000 - 2600$; this parameter largely sets the "range" of our fold-change vs fold-change curves and this range of $\alpha$ reproduces the experimental range we see in those curves for this data set. We then use this same on rate to derive the relevant off rates for O2 and Oid using their binding energies ($\epsilon_{\mathrm{O2}} = 13.9$ $k_{\mathrm{B}}T$ , $\epsilon_{\mathrm{Oid}} = 16.3\ k_{\mathrm{B}}T$) and Eqn. S1. Interestingly, the binding affinity we measure for Oid is $0.7 k_{\mathrm{B}}T$ weaker than has been previously reported but is consistent with measurements of Oid binding affinity in our lab. Using this method, we find the $k_{\mathrm{on}}$ to be 0.0015 per TF per second, which yields $k_{\mathrm{off}}$ to be, O1=0.0015 $s^{-1}$, O2 = 0.0167 $s^{-1}$ and Oid=0.0004 $s^{-1}$, consistent with previous findings (*27,31,41,44*) All other rates are listed in Table **S2**.

Importantly, this process is not meant to precisely determine the exact quantitative parameters of LacI binding, and it is not a formal fit, but rather an estimate that provides us with realistic prediction of regulation from our simulations using molecular parameters that are consistent with available direct kinetic measurements (*37,72-74*).

#### Data Analysis

Data analysis is performed using a modified version of the Matlab code Schnitzcells (*51*). We use this code to segment the phase images of each sample to identify single cells. Mean pixel intensities of YFP and mCherry signals are extracted from the segmented phase mask for each individual cell using regionprops, an inbuilt function in matlab. The background fluorescence is calculated by averaging the mean intensity of the inverse phase mask upon eroding the regions around the segmented cell masks. The background fluorescence value of a particular frame was subtracted from the mean pixel intensity of cells in the same frame (see SI text **S1**). Finally, the autofluorescence value were calculated using the same procedure for cells that do not express either YFP or mCherry and the average autofluorescence value of these cells is subtracted from each measured YFP or mCherry value. Resulting mean pixel intensity of mCherry signal was corrected for the crosstalk from YFP signal. Crosstalk between different channels can be measured by determining the difference between the autofluorescence of a strain without a given fluorophore in the presence of the other fluorophore (highly expressed). We find that under our microscope $0.25\%$ ($\gamma_{\mathrm{cross}}$=0.0025) of YFP signals can be seen in the mCherry channel whereas mCherry channel has no crosstalk in the YFP channel. Hence, we correct for this crosstalk by subtracting the mean pixel intensity of YFP signal times the $\gamma_{\mathrm{cross}}$ from the mean pixel intensity of mCherry signal. The per pixel fluorescence values of mCherry and YFP of each cell is then multiplied by the area of the cell to account for the total fluorescence. Fold-change in expression of the mCherry and YFP are calculated by dividing the corresponding values of the constitutive strains (discussed in SI text **S4**). At least 500 individual cells were analyzed per sample and binned according to the mCherry values.

#### Supplementary Text

##### S1. Sensitivity in choosing the background values

The local background of each image is subtracted from individual cells of that image, rather than using a global average over every position. Getting a precise quantitative measurement of fluorescence values is important especially for the tagged strains as their mCherry signal can be only several counts above autofluorescence. The background fluorescence can be influenced by factors such as the local thickness of the agarose pad and positional effects due to the glass dish (which can have small local defects). As shown in SI Fig. **S1A-B**, a no fluorescent strain corrected using the local fluorescence (calculated by making an inverse mask of each frame, excluding regions with cell, and calculating the mean intensity of the background) of each frame produces a tight, symmetric distribution of cell fluorescence with the mean centered near 0 when compared to using the mean value of no fluorescent strain. In other words, many of the YFP or mCherry signals that appear high in the autofluorescence samples also have higher than average backgrounds and thus accounting for this image to image difference is important. Hence, for all experiments we have used the local background fluorescence of each frame to correct for the autofluorescence of cells in the corresponding frame and excluding frames with too high variation in the background fluorescence.

##### S2. Cell growth rate in different media

Cell growth rate is measured in strain HG105 growing in a 50 mL flask at 37°C and at 250 rpm. Samples are collected at precise time points and OD600 is measured. Doubling time is calculated by first interpolating the intermediate time points from the measurements of OD600 and with the single exponential robust fit function in matlab (see SI Fig. **S2A**). SI Fig. **S2B** shows the scaling in cell area (measured in pixel units) in different media in accordance with the previous literature (*75*). Interestingly, the strains with 4X and 5X decoys have a strikingly different area (from other strains) possibly indicating sickness due to the presence of multiple arrays of Oid binding site.

##### S3. Quantification of plasmid copy number

Five different variants of Oid decoy arrays (carrying 1, 2, 3, 4 and 5 binding sites for Oid, respectively) are inserted in the intergenic region between the origin of replication and ampicillin cassette of the pZE plasmid. Plasmid copy number is quantified in qPCR measurements using primers that targets a 90 bp-intergenic region immediately upstream of the site of insertion of our decoy array. The total number of decoys can then be calculated by multiplying the measured copy number of pZE plasmid with the number of binding sites in the decoy array. As shown in SI Fig. **S3A**, pZE plasmid copy number is similar in strains with different decoy arrays except for strains carrying the 5X decoy array plasmid. Copy number of 5X-decoy array plasmid is significantly higher when compared to strains carrying other decoy array plasmids. This difference is primarily due to a reduced CFU/mL obtained (see SI Fig. **S3C**) for strains carrying the 5X decoy arrays; the number of molecules of plasmid per reaction is uniform across different strains (see SI Fig.

**S3B**). It is not clear if this is due to this sample actually containing less cells or if it is due to a reduced ability to recover and separate these cells (which tend to clump and stick more in microscopy imaging) in the plating assay. This may lead to over-prediction of the copy number of 5X decoy plasmid. Hence, we excluded the copy number of 5X-decoy plasmid and estimated the average copy number of pZE plasmid to be 48 copies/cell (SI Fig. **S3A**). The average ($\pm$ standard deviation) number of decoy binding arrays in different strains are: 39$\pm$8, 96$\pm$17, 134$\pm$25, 245$\pm$40, and 607$\pm$47, respectively.

#### S4. Constitutive values for the autoregulatory gene

To compare expression levels between the TF and the target genes, we wish to compare fold-change as an "apples-to-apples" comparison of the regulation of each gene. To calculate fold-change we must know the constitutive expression of the gene, *i.e.* how much expression is seen in the absence of regulation by TF. In simulation, this is simple to calculate because we can remove any reactions that include TF binding. Experimentally, calculating constitutive expression for the target gene is also relatively straight-forward; we delete the gene expressing LacI-mCherry and measure the same construct in the absence of TF. However, measuring constitutive expression experimentally for an autoregulating gene was more challenging. There are many possible strategies, but all of them come with some complication. In short, we attempted 3 different strategies which included: (1) IPTG induction (with or without the addition of decoys), (2) mutated LacI to ablate specific binding, (3) mutated binding site sequences (which has the complication that the site is centered at +11 and thus is both close to the promoter and present on the transcript, see SI Fig. **S4A**). In the end, we identified one mutated site (NoO1V1) which faithfully preserved constitutive expression of the target gene in all media studied. Unfortunately, we were not able to find corresponding mutated sites that reproduced expression of promoters bearing O2 or Oid binding sites. As such, for data using those binding sites on the TF gene we have an unknown scaling factor between the x- and y-axis in the fold-change versus fold-change plots which we determine by fitting the glucose data to our simulations (and then hold constant for all other data sets). In the following sections we discuss techniques we tried.

##### *S4.1 Allosteric induction with IPTG to achieve constitutive expression*

One way to obtain the constitutive values is to exploit the property of the LacI to become less active when bound to small molecules like IPTG. Previous studies indicate that even with the use of IPTG, expression from a stronger binding site (like Oid) cannot be fully rescued when the repressor copy number is high (*76*). In our experiments, we observed this phenomenon as well. We thought that perhaps if we also had a large number of decoy binding sites present we could recover full expression of the TF gene. As shown in SI Fig. **S4C-E**, for most strains expressing the TF, the expression of the target could not be fully rescued with IPTG and decoys. Hence, allosteric induction with IPTG could not serve as a right constitutive value for our system.

##### *S4.2. Use of LacI with mutated DNA recognition domains*

We constructed a mutant protein by deleting 10 amino acids (from amino acid 60 to amino acid 70) in the DNA binding domain of LacI. This mutant helped to completely restore the target expression. However, fold-change in mCherry fluorescence was greater than one for lowly expressed TF levels. This discrepancy may originate from many possible sources such as a change to the stability of the mRNA/protein or a possible alteration to the spectral property of mCherry (which is directly fused to LacI). In the end we were unable to find a suitable LacI mutant without this feature.

##### *S4.3. Use of binding sequence insensitive to LacI*

Oehler *et al.* 1994 has reported inactivated O1 site (NoO1V1) that has close consensus to O1 binding sequence but does not allow LacI binding. We verified that the expression of YFP from the promoter with NoO1V1 is comparable to the expression of YFP from O1 regulated promoter (in the absence of any LacI) but is lower than the expression from O2 and Oid regulated promoters (SI Fig. **S4B**). Although expression alone does not guarantee that all intermediate steps are precisely the same, we believe this construct gives accurate measurements of constitutive expression for the TF and target genes. We used TF and target with NoO1V1 binding sequence as our constitutive strain to normalize expression from any O1 regulated genes in our experiments. We also tried other forms of mutations on the NoO1V1 binding site (SI Fig. **S4A**) in order to obtain mutants that relives *lacI* repression and restore expression of Oid or O2 sequence but with no success.

#### S5. Copy number difference and Diffusion limitation of TF

Copy number variation of genes along the long axis of the chromosome and the diffusion limitation of LacI-mCherry could be suggested as a significant contributor to the asymmetry between TF and the target. *E. coli* can initiate multiple replication events (depending on the division rate in the given media) and hence different genes along the chromosome will experience a different copy number in a given time. For instance, *E. coli* growing in RDM (with a division rate of 22 minutes) will have a copy number of 4 at the *ybcN* locus (where the TF gene is integrated) and a copy number of 3.6 at the *galK* locus (where the target gene is integrated) as described by Cooper *et al.* (*77*). We believe that the use of fold-change as the measurement of expression helps to reduce the influence of copy number effects (since both the regulated and unregulated measurements have the same copy number). However, the effects may not be linear and LacI has been shown to suffer from diffusion limitation from its origin of synthesis (*78*). Hence, we tested our system by placing the TF and the target genes integrated next to each other at the *gspI* locus. As evident from SI Fig. **S5**, there is no significant contribution of the copy number difference between TF and target or diffusion limitation of TF on the phenomenon of asymmetry observed in our negatively-autoregulated SIM motif.

#### S6. Deterministic solution

Using the assumptions of equilibrium mass-action kinetics, the deterministic counterpart of the negative autoregulation system described above can be written as

$$
\begin{aligned}
\frac{dX}{dt} &= \alpha m_{\mathrm{x}} - \gamma X + 2k_{\mathrm{m}}Z - 2k_{\mathrm{p}}X^2, \\
\frac{dZ}{dt} &= -k_{\mathrm{m}}Z + k_{\mathrm{p}}X^2 - \gamma Z - k_{\mathrm{on}}ZP_{\mathrm{fx}} - k_{\mathrm{on}}ZP_{\mathrm{fy}} - k_{\mathrm{on}}ZN_{\mathrm{f}} + k_{\mathrm{off,x}}(1 - P_{\mathrm{fx}}) \\
&\qquad + k_{\mathrm{off,y}}(1 - P_{\mathrm{fy}}) + k_{\mathrm{off,d}}(N - N_{\mathrm{f}}), \\
\frac{dY}{dt} &= \alpha m_{\mathrm{y}} - \gamma Y, \\
\frac{dP_{\mathrm{fx}}}{dt} &= -k_{\mathrm{on}}ZP_{\mathrm{fx}} + (k_{\mathrm{off,x}} + \gamma)(1 - P_{\mathrm{fx}}), \\
\frac{dP_{\mathrm{fy}}}{dt} &= -k_{\mathrm{on}}ZP_{\mathrm{fy}} + (k_{\mathrm{off,y}} + \gamma)(1 - P_{\mathrm{fy}}), \\
\frac{dN_{\mathrm{f}}}{dt} &= -k_{\mathrm{on}}ZN_{f} + (k_{\mathrm{off,d}} + \gamma)(N - N_{\mathrm{f}}), \\
\frac{dm_{\mathrm{x}}}{dt} &= \beta P_{\mathrm{fx}} - \gamma_m m_{\mathrm{x}}, \\
\frac{dm_{\mathrm{y}}}{dt} &= \beta P_{\mathrm{fy}} - \gamma_m m_{\mathrm{y}}.
\end{aligned} \tag{S2}
$$

Here, $X$ is the concentration of free TF monomer, $Y$ is the concentration of target protein, and $Z$ is the concentration of TF dimer. $m_{\mathrm{x}}, m_{\mathrm{y}}, P_{\mathrm{fx}}(P_{\mathrm{ox}}), P_{\mathrm{fy}}(P_{\mathrm{oy}}), N$, and $N_{\mathrm{f}}(N_{\mathrm{o}})$ are TF mRNA, target mRNA, free (bound) TF-promoter, free (bound) target-promoter, total concentration of decoy sites, and concentration of free (bound) decoy sites, respectively. Inherent in the equations are the assumptions of the conservation for the concentration of binding sites , i.e. $P_{\mathrm{fx}} + P_{\mathrm{ox}} = 1$, $P_{\mathrm{fy}} + P_{\mathrm{oy}} = 1$, and $N_{\mathrm{f}} + N_{\mathrm{o}} = N$. The right hand side of the equations can be set to zero to obtain the steady state values for all the components.

$$
\begin{aligned}
P_{\mathrm{fx}} &= \frac{k_{\mathrm{off,x}} + \gamma}{k_{\mathrm{on}}Z + k_{\mathrm{off,x}} + \gamma} = \frac{1}{1 + \sigma_1 Z}, \\
P_{\mathrm{fy}} &= \frac{k_{\mathrm{off,y}} + \gamma}{k_{\mathrm{on}}Z + k_{\mathrm{off,y}} + \gamma} = \frac{1}{1 + \sigma_2 Z}, \\
N_{\mathrm{f}} &= \frac{N(k_{\mathrm{off,d}} + \gamma)}{k_{\mathrm{on}}Z + k_{\mathrm{off,d}} + \gamma} = \frac{N}{1 + \sigma_3 Z}, \\
m_{\mathrm{x}} &= \frac{\beta}{\gamma_{\mathrm{m}}}P_{\mathrm{fx}}, \\
m_{\mathrm{y}} &= \frac{\beta}{\gamma_{\mathrm{m}}}P_{\mathrm{fy}}, \\
0 &= \alpha m_{\mathrm{x}} - \gamma X + 2k_{\mathrm{m}}Z - 2k_{\mathrm{p}}X^2, \\
0 &= -k_{\mathrm{m}}Z + k_{\mathrm{p}}X^2 - \gamma(Z + P_{\mathrm{ox}} + P_{\mathrm{oy}} + N_{\mathrm{o}} \\
Y &= \frac{\alpha\beta}{\gamma\gamma_{\mathrm{m}}}P_{\mathrm{fy}} = \frac{\alpha\beta}{\gamma\gamma_{\mathrm{m}}}\frac{1}{1 + \sigma_1 X},
\end{aligned} \tag{S3}
$$

where $\sigma_i = k_{\text{on}}/(k_{\text{off,i}} + \gamma)$. The concentration of total TF protein can be expressed as a sum of free TF monomer, TF dimer bound to each promoter, and TF dimers bound to the decoys sites

$$
\begin{aligned}
X_{\text{Total}} &= X + 2(Z + P_{\text{ox}} + P_{\text{oy}} + N_{\text{o}}), \\
&= \frac{\alpha}{\gamma} m_{\text{x}}, \\
&= \frac{\alpha\beta}{\gamma\gamma_{\text{m}}} \frac{1}{1+\sigma_1 Z}. \qquad \text{(S4)}
\end{aligned}
$$

The fold-change of the TF and target expression, thus can be obtained by dividing $X_{\text{Total}}$ and $Y$ with the constitutive expression, i.e., $\alpha\beta/\gamma\gamma_{\text{m}}$ which yields,

$$
\begin{aligned}
\text{FC}_{\text{TF}} &= \frac{1}{1+\sigma_1 Z} = \frac{1}{1+\frac{k_{\text{on}}}{k_{\text{off,x}}+\gamma} Z}, \qquad \text{(S5)} \\
\text{FC}_{\text{Target}} &= \frac{1}{1+\sigma_2 Z} = \frac{1}{1+\frac{k_{\text{on}}}{k_{\text{off,y}}+\gamma} Z}. \qquad \text{(S6)}
\end{aligned}
$$

It is worth noting that both TF and target protein follows $1/(1+R^*)$, where $R^* = Zk_{\text{b}}/(k_{\text{u}}+\gamma)$ is the reduced free TF concentration, which is equivalent to the thermodynamic solution (48). When the unbinding rates of TF and target are identical, each of them follow the same fold-change curve irrespective of the competition from other decoy sites. In SI Fig. **S**7C, we plot the fold-change for TF and target with $k_{\text{off,x}}$ corresponding to O1 binding site and $k_{\text{off,y}}$ corresponding to O1 (yellow), O2 (purple), and Oid (blue). It can be seen from the figure that when the off-rates are identical the fold-change curve follows one-to-one line showing no asymmetry which is in contrast with the results obtained using stochastic simulations and experimental results. Furthermore, both the transient and steady state behavior of mean fold-change of TF and target obtained from deterministic solution deviate from the stochastic behavior (see SI Fig. **S**7B). Importantly, when autoregulation is removed from the simulation, the deterministic and stochastic solutions agree precisely (SI Fig. **S**7B inset).

#### S7. Maximum asymmetry

The asymmetry in regulation (defined as $\text{FC}_{\text{TF}} - \text{FC}_{\text{Target}}$) is a function of all the rates describing the system and number of decoy binding sites. For a given set of rates ($k_{\text{on}}$, $k_{\text{off}}$, $\gamma, \gamma_m$) as the decoy number is varied the asymmetry first increases, attains a maximum and then approaches zero for infinite number of decoy binding sites (see SI Fig. **S**8). The maximum asymmetry for a given set of rates is this peak asymmetry observed as decoy number is varied. In the manuscript we show a heatmap (Fig. **3**E) to emphasize how this maximum asymmetry depends on the two crucial rate parameters, off-rate of the binding sites ($k_{\text{off}}$ or equivalently binding affinity, since in our model $k_{\text{on}}$ is kept constant) and the degradation of TF molecules ($\gamma$).

#### S8. A minimal model of an autoregulatory gene and a single target gene

The full model of section SI text **S6** contains many reactions that are included to more faithfully mirror the biological system we are modeling. However, not all of these reactions are necessary to observe the phenomenon of asymmetry which we describe in this manuscript. In this section, we present a reduced model of the extended model of transcription described in Materials and methods to show that the asymmetry in TF and target expression stems from the network architecture and not due to the intermediate steps of transcription and the presence of excess decoy binding sites. We consider an autoregulatory gene whose protein product $X$ inhibits its own expression and also represses a single target gene with protein product $Y$. To reduce the complexity, the protein is made directly from the gene with no intermediates (eliminating translation rates and mRNA decay rates). In this system the TF, $X$, acts as a monomer and binds to its own gene with rate $k_{\text{on}}$ and unbinds with rate $k_{\text{off}}$. Similarly, the TF ($X$) binds and unbinds from the target gene with the same rates. Both the TF gene and target gene in free state (not bound with TF) produces their protein with rate $\alpha$ which degrades with rate $\gamma$ (dilution through cell division). The reactions describing this reduced model are listed in Table **S1B**. We implement the simulations using stochastic simulation algorithms as described in Materials and Methods section.

Next, we write a set of deterministic coupled ODEs corresponding to the reactions described above which is given by

$$
\begin{aligned}
\frac{dX}{dt} &= \alpha P_{\text{fx}} - \gamma X - k_{\text{on}} X P_{\text{fx}} - k_{\text{on}} X P_{\text{fy}} + k_{\text{off}}(1 - P_{\text{fx}}) + k_{\text{off}}(1 - P_{\text{fy}}), \\
\frac{dY}{dt} &= \alpha P_{\text{fy}} - \gamma Y, \\
\frac{dP_{\text{fx}}}{dt} &= -k_{\text{on}} X P_{\text{fx}} + (k_{\text{off}} + \gamma)(1 - P_{\text{fx}}), \\
\frac{dP_{\text{fy}}}{dt} &= -k_{\text{on}} X P_{\text{fy}} + (k_{\text{off}} + \gamma)(1 - P_{\text{fy}}),
\end{aligned}
\tag{S7}
$$

Here, $X$ is the concentration of free TF and $Y$ is the concentration of target protein. $P_{\text{fx}}(P_{\text{ox}})$ and $P_{\text{fy}}(P_{\text{oy}})$ are free (bound) TF-promoter, free (bound) target-promoter, respectively. Inherent in the equations are the assumptions of the conservation for the concentration of binding sites, i.e. $P_{\text{fx}} + P_{\text{ox}} = 1$, $P_{\text{fy}} + P_{\text{oy}} = 1$. To obtain the steady state values of TF and target expression the right hand side of the equations is set to zero which yield

$$
\begin{aligned}
P_{\text{fx}} &= \frac{k_{\text{off}} + \gamma}{k_{\text{on}} X + k_{\text{off}} + \gamma} = \frac{1}{1 + \sigma X}, \\
P_{\text{fy}} &= \frac{k_{\text{off}} + \gamma}{k_{\text{on}} X + k_{\text{off}} + \gamma} = \frac{1}{1 + \sigma X}, \\
X &= \frac{\alpha}{\gamma} P_{\text{fx}} - P_{\text{ox}} - P_{\text{oy}}, \\
Y &= \frac{\alpha}{\gamma} P_{\text{fy}} = \frac{\alpha}{\gamma} \frac{1}{1 + \sigma X},
\end{aligned}
\tag{S8}
$$

where $\sigma = k_{\rm on}/(k_{\rm off} + \gamma)$. Total TF concentration, $X_{\rm Total}$, can be expressed as the sum of free TF and TFs bound to each promoter

$$\begin{aligned} X_{\rm Total} &= X + P_{\rm ox} + P_{\rm oy} \\ &= \frac{\alpha}{\gamma} P_{\rm fx} \\ &= \frac{\alpha}{\gamma} \frac{1}{1 + \sigma X}. \end{aligned} \tag{S9}$$

The fold-change of the TF and target expression, thus can be obtained by dividing $X_{\rm Total}$ and $Y$ by the constitutive expression, *i.e.* without any regulation, $C_0 = \alpha/\gamma$ which yields,

$$\mathrm{FC}_{\rm TF} = \frac{1}{1+\sigma X} = \frac{1}{1+\frac{k_{\rm on}}{k_{\rm off}+\gamma} X}, \tag{S10}$$

$$\mathrm{FC}_{\rm Target} = \frac{1}{1+\sigma X} = \frac{1}{1+\frac{k_{\rm on}}{k_{\rm off}+\gamma} X}. \tag{S11}$$

As was shown previously in section **S6**, both TF and target protein follows $1/(1 + \sigma X)$ and show no asymmetry in regulation.

Furthermore, solving Eqn. S8 we get the free TF expression as,

$$X = \frac{-1 - 2\sigma + \sqrt{(1 + 2\sigma)^2 + 4C_0\sigma}}{2\sigma}, \tag{S12}$$

In Fig. **S**9 we plot the asymmtery as a function of growth rate (inverse of degradation) obtained from stochastic simulation and the solution from deterministic ODE.

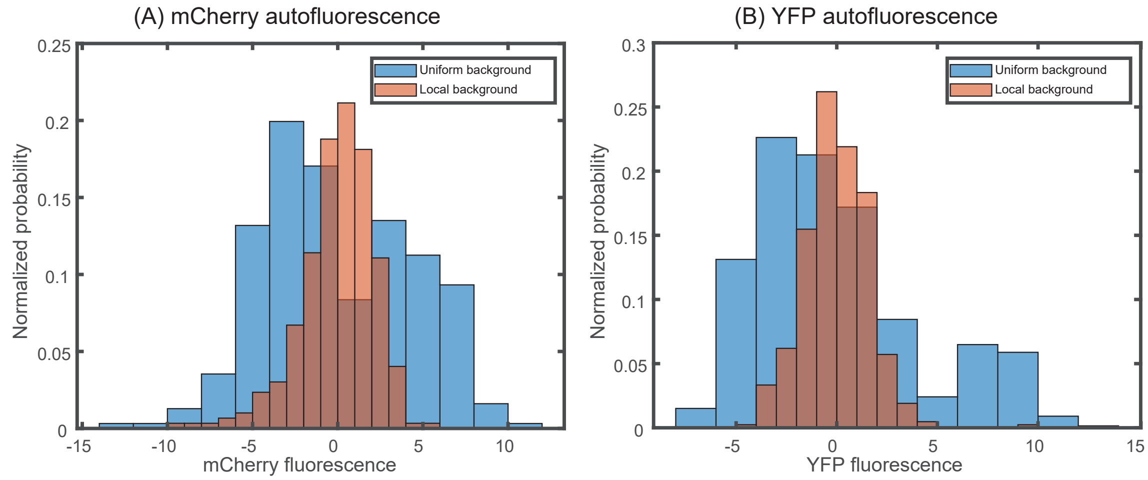


Figure S1: **Accounting for local variation in background fluorescence**. Histogram of single-cell autofluorescence levels of (**A**) mCherry or (**B**) YFP fluorescence in a strain without the YFP and mCherry casettes. The blue bars are calculated as the fluorescence level subtracted from the average across the entire sample (9 different fields of view). The red bars are calculated by first removing the local background fluorescence from cells at each position before subtracting the remaining signal from the average. The wide distribution seen in the blue bars is owed largely to local differences in background fluorescence and is removed by accounting for position-to-position variability.

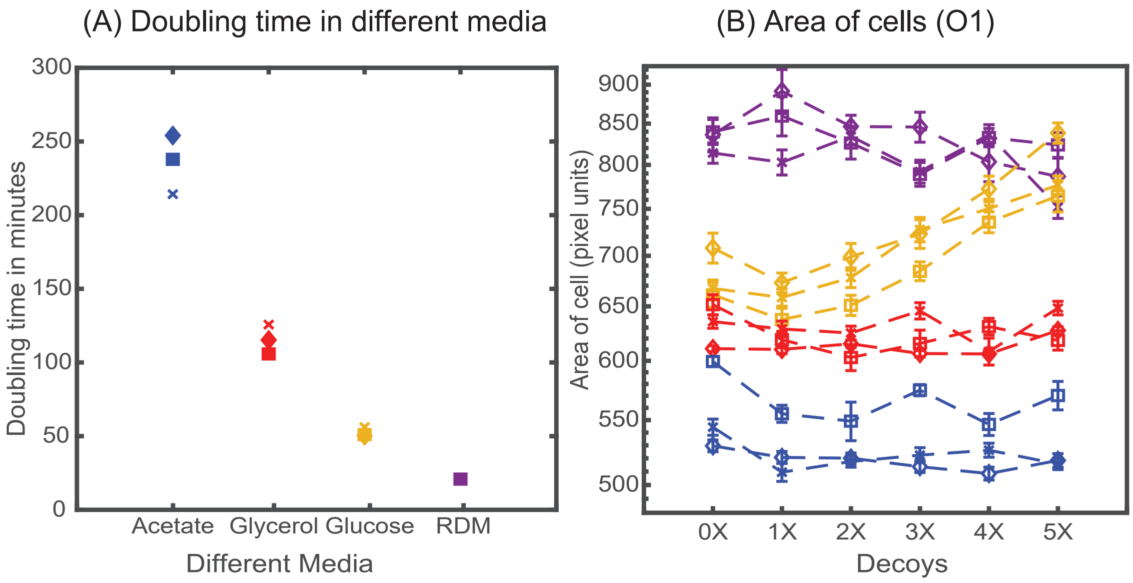


Figure S2: **Growth rate in different media**. (**A**) Doubling time of HG105 in different media used in this study. (**B**) Consistent with the literature there is a scaling of cell area in different media in accordance with their growth rate. Strains with 4X and 5X decoys growing in glucose minimal media have a drastically different cell area.

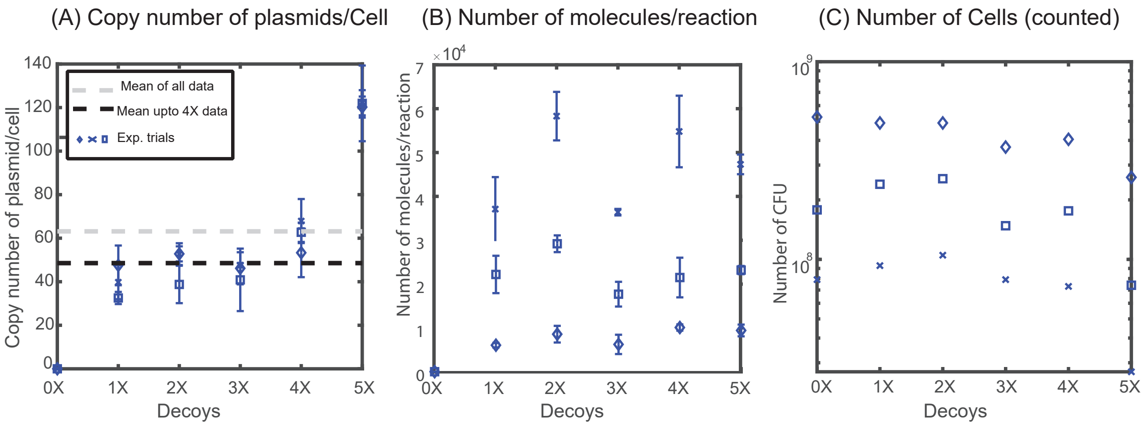


Figure S3: **Quantification of plasmid copy number**. (**A**) Copy number of decoy array plasmids measured in M9-Glucose minimal media. (**B**) Number of molecules obtained per qPCR reaction remains constant across different decoy strains (1X, 2X, 3X, 4X, 5X). (**C**) Number of Colony Forming Units (CFU) per mL used to normalize the number of molecules to account for the copy number of plasmids per cell.

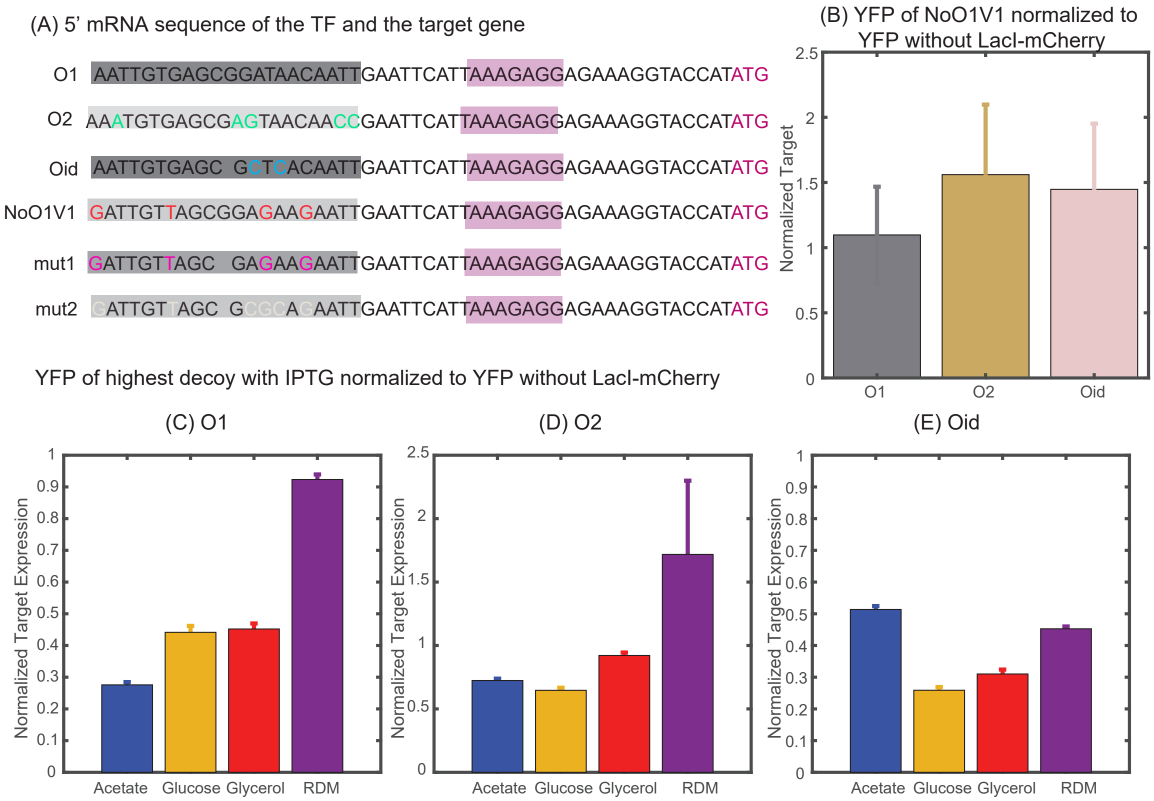


Figure S4: **Determining constitutive expression of YFP and mCherry**. (**A**) 5′ mRNA sequence of the TF and the target genes. The binding site for the TF is carried in the mRNA sequence and is highlighted in shaded dark grey boxes with base changes for different binding sites coded in multicolor. mut1 and mut2 are the two variant binding sites that are designed with mutations similar to NoO1V1 but with Oid site length. However, such changes do not achieve constitutive unregulated expression similar to O2 or Oid. (**B**) Plot showing YFP expressed from NoO1V1 regulated promoter normalized to YFP expressed from promoter regulated with O1, O2 or Oid. (**C-E**) Plot showing the effect of IPTG in relieving YFP expression form O1 (**C**), O2 (**D**) or Oid. (**E**) regulated promoter and with 5X decoy plasmids. As indicated in the plot IPTG is not sufficient to restore complete expression of YFP in different media and hence cannot be used as a measure of constitutive expression.

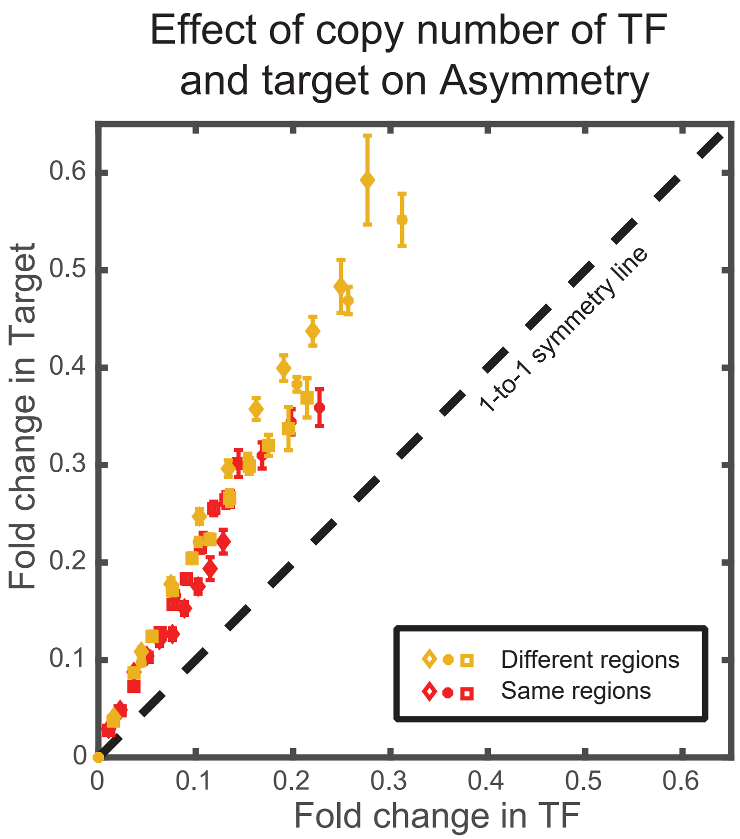


Figure S5: **Effect of copy number difference on asymmetry**. Comparison of asymmetry in strain where the TF and the target genes are located either at two different regions of the chromosome (*ybcN* for TF and *galK* for target, shown in yellow data points)) or when it is present together in the chromosome (at the *gspI* locus, shown in red data points).

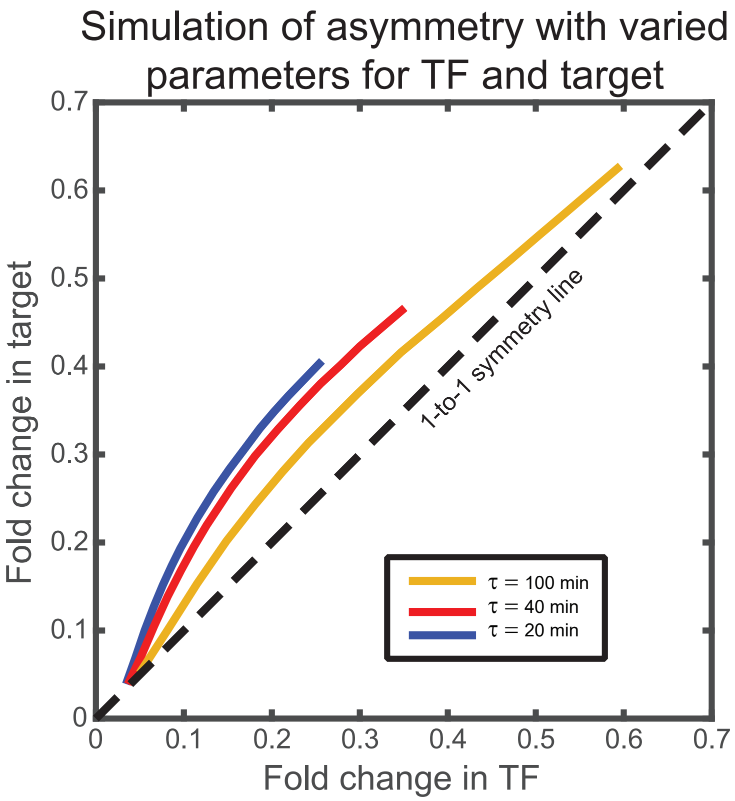


Figure S6: **Asymmetry for different growth rate ($\tau$) with varying transcription rate, translation rate, and mRNA stability**. Stochastic simulation performed using the kinetic parameters listed in Bremer and Dennis (*42*) for $\tau$ being 20 (blue line), 40 (red line), and 100 (yellow line) minutes. The qualitative ordering and features of the asymmetry curve is not impacted by the changes in the kinetic parameters such as transcription rate, translation rate, and mRNA stability due to change in growth rates.

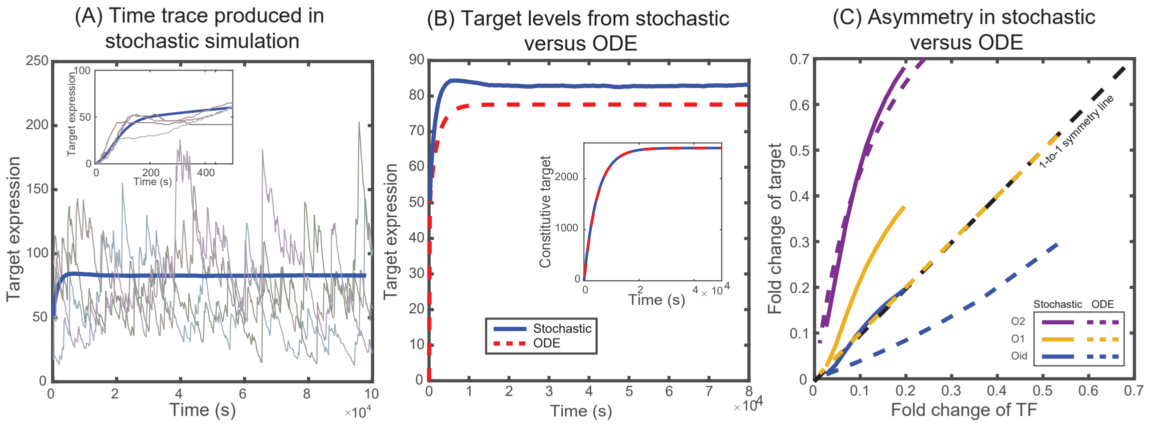


Figure S7: **Solutions from stochastic simulation and from deterministic ODEs**. (**A**) Representative time traces of target expression in individual cells (grey shades) from stochastic simulations. Blue solid line represents the mean behavior averaged over $5 \times 10^4$ iterations. Inset shows the transient behavior. (**B**) Plot showing the average target expression in the negative SIM motif from stochastic simulations (solid line) and from solving deterministic ODEs (dashed line). Inset shows that when regulation is removed the average levels are identical for stochastic and deterministic models. (**C**) Plot showing the asymmetry between TF and target expression from using either stochastic simulation (solid lines) or solving deterministic ODEs (dashed lines). The TF is always regulated by O1 binding site whereas the target is regulated by O1 (yellow), O2 (purple) or Oid (blue) binding sites. The black dashed line represents line of no asymmetry.

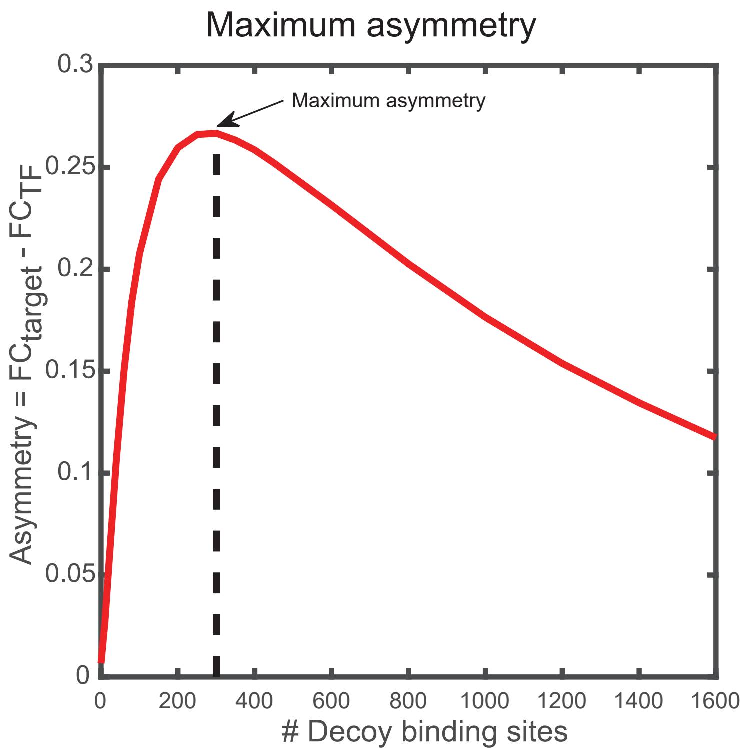


Figure S8: **Determination of maximum asymmetry**. Maximum asymmetry in simulation is computed by plotting the asymmetry, difference in fold-change between target and TF, versus number of decoy binding sites in SIM motif. The peak of this asymmetry corresponds to the maximum asymmetry.

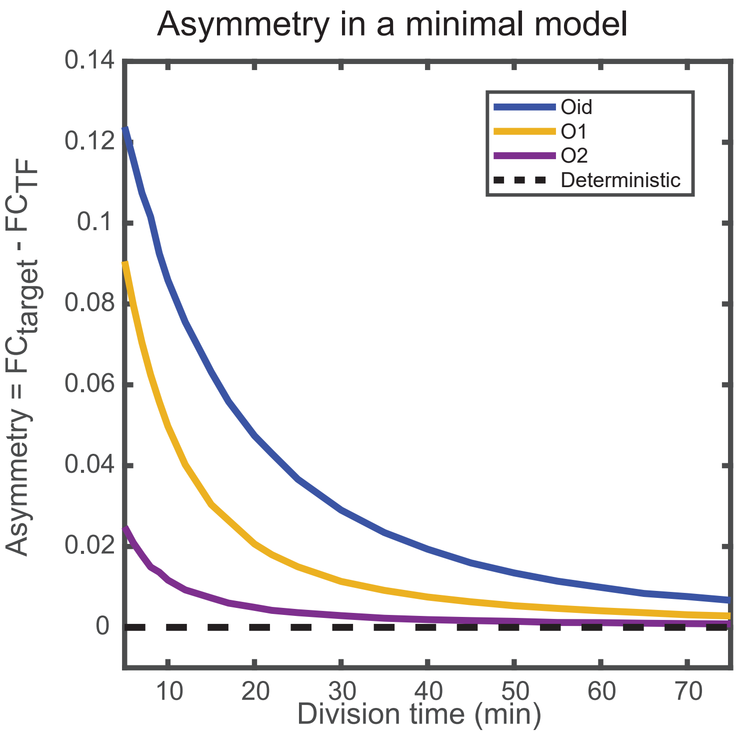


Figure S9: **Minimal model of autoregulation**. Asymmetry predicted from a minimal model without intermediate transcription steps and decoy binding sites. The asymmetry follows similar trend as predicted in the complete stochastic model. Stronger binding site (Oid, shown in solid blue line) shows higher asymmetry than a weak binding site (O2, shown in solid purple line). Also, asymmetry decreases as the growth rate is increased. Dashed line corresponds to the deterministic counterpart of the stochastic reaction systems. Again, we do not find any asymmetry in TF and target regulation from the deterministic solution.

Table S1: **List of reactions used in the (A) stochastic model and (B) in the minimal model.**

#### (A) Full model

| Schematic reaction | Stochastic chemical reaction | Deterministic rate |
|---|---|---|
| 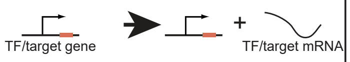  | $m_{\mathrm{TF(Target)}} \xrightarrow{\beta} m_{\mathrm{TF(Target)}} + 1$ | $\beta P_{\mathrm{fx(fy)}}$ |
| 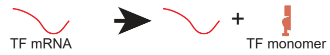  | $m_{\mathrm{TF}} \xrightarrow{\alpha} m_{\mathrm{TF}} + X$ | $\alpha m_{\mathrm{x}}$ |
| 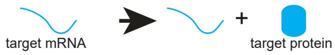  | $m_{\mathrm{Target}} \xrightarrow{\alpha} m_{\mathrm{Target}} + Y$ | $\alpha m_{\mathrm{y}}$ |
| 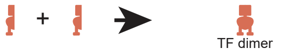  | $X + X \xrightarrow{k_{\mathrm{p}}} Z$ | $k_{\mathrm{p}} X^2$ |
| 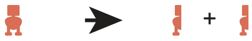 | $Z \xrightarrow{k_{\mathrm{m}}} X + X$ | $k_{\mathrm{m}} Z$ |
| 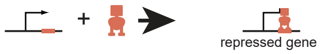  | $Z + P_{\mathrm{TF(Target)}} \xrightarrow{k_{\mathrm{on}}} ZP_{\mathrm{TF(Target)}}$ | $k_{\mathrm{on}} Z P_{\mathrm{fx(fy)}}$ |
| 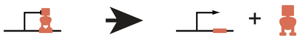 | $ZP_{\mathrm{TF(Target)}} \xrightarrow{k_{\mathrm{off,TF(Target)}}} Z + P_{\mathrm{TF(Target)}}$ | $k_{\mathrm{off,x(off,\ y)}}(1 - P_{\mathrm{fx(fy)}})$ |
| 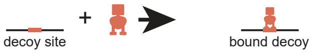  | $Z + N \xrightarrow{k_{\mathrm{on}}} ZN$ | $k_{\mathrm{on}} Z N_f$ |
| 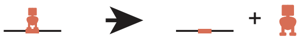 | $ZN \xrightarrow{k_{\mathrm{off,Decoy}}} Z + N$ | $k_{\mathrm{off,d}} (N - N_{\mathrm{f}})$ |
| 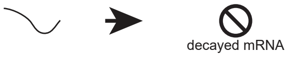  | $m_{\mathrm{TF(Target)}} \xrightarrow{\gamma_{\mathrm{m}}} m_{\mathrm{TF(Target)}}$ | $\gamma_m m_{\mathrm{x(y)}}$ |
| 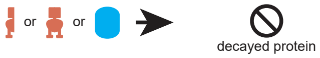  | $X, Y, Z \xrightarrow{\gamma} \phi$ | $\gamma X$ or $\gamma Y$ or $\gamma Z$ |
| 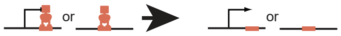  | $ZP_{\mathrm{TF(Target)}} \xrightarrow{\gamma} P_{\mathrm{TF(Target)}}$ or $ZN \xrightarrow{\gamma} N$ | $\gamma(1 - P_{\mathrm{fx(fy)}})$ or $\gamma(N - N_{\mathrm{f}})$ |

#### (B) Minimal model to demonstrate asymmetry

| Schematic reaction | Stochastic chemical reaction | Deterministic rate |
|---|---|---|
| 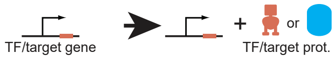  | $P_{\mathrm{TF(Target)}} \xrightarrow{\alpha} P_{\mathrm{TF(Target)}} + X(Y)$ | $\alpha P_{\mathrm{fx(fy)}}$ |
| 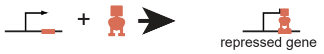  | $X + P_{\mathrm{TF(Target)}} \xrightarrow{k_{\mathrm{on}}} XP_{\mathrm{TF(Target)}}$ | $k_{\mathrm{on}} X P_{\mathrm{fx(fy)}}$ |
| 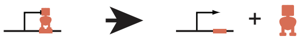 | $XP_{\mathrm{TF(Target)}} \xrightarrow{k_{\mathrm{off}}} Z + P_{\mathrm{TF(Target)}}$ | $k_{\mathrm{off}}(1 - P_{\mathrm{fx(fy)}})$ |
| 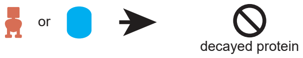  | $X, Y \xrightarrow{\gamma} \phi$ | $\gamma X$ or $\gamma Y$ |
| 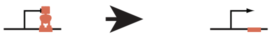 | $XP_{\mathrm{TF(Target)}} \xrightarrow{\gamma} P_{\mathrm{TF(Target)}}$ | $\gamma(1 - P_{\mathrm{fx(fy)}})$ |

Table S2: **Kinetic rates used in the simulations**

| Rates | Symbols | Value | Reference |
|---|---|---|---|
| Growth rate | $1/\gamma$ | 25 min (RDM),<br>55 min (Glucose),<br>125 min (Glycerol),<br>225 min (Acetate) | Measured experimentally |
| Binding of TF | $k_{\mathrm{on}}$ | 0.0015 $\mathrm{TF}^{-1}\mathrm{s}^{-1}$ | Obtained from fit |
| Unbinding of TF | $k_{\mathrm{off}}$ | 0.00042 $\mathrm{s}^{-1}$ (Oid)<br>0.00149 $\mathrm{s}^{-1}$ (O1)<br>0.0167 $\mathrm{s}^{-1}$ (O2) | Eqn. S1 |
| mRNA degradation | $\gamma_m$ | 0.033 $\mathrm{s}^{-1}$ | Obtained from fit |
| mRNA production | $\beta$ | 0.1 $\mathrm{s}^{-1}$ | Obtained from fit |
| Translation rate | $\alpha$ | 0.033 $\mathrm{s}^{-1}$ | Obtained from fit |
| Dimerization | $k_{\mathrm{p}}$ | 1.38 $\mathrm{s}^{-1}$ | (*79*) |
| Monomerization | $k_{\mathrm{m}}$ | 0.000002 $\mathrm{s}^{-1}$ | (*79*) |

Table S3: **Primers used in this study are listed below.** Primers for the chromosomal integration of TF and the target are the same as described in (*23*). Primers to mutate the binding sites from O1 to Oid, O2 or NoO1V1 is listed below with the binding sites highlighted in yellow. Primers to introduce the degradation tags to LacI mCherry fusion protein is listed below with tag sequence represented in red.

| **Mutagenesis Primer** | |
|---|---|
| Oid_ FP | CCGGCTCGTATAATGTGTGGAATTGTGAGCGCTCACAATTGAATTCATTAAAGAG |
| Oid_ RP | CTCTTTAATGAATTCAATTGTGAGCGCTCACAATTCCACACATTATACGAGCCGG |
| O2_ FP | GTGAGCGAGTAACAACCGAATTCATTAAAGAGGAGAAAGGTAC |
| O2_ RP | TTGTTACTCGCTCACATTTCCACACATTATACGAGCC |
| NoO1V1_FP | GATTGTTAGCGGAGAAGAATTGAATTCATTAAAGAGGAGAAAGGTACC |
| NoO1V1_RP | AATTCTTCTCCGCTAACAATCCCACACATTATACGAGCCGGAAG |
| **Primers to introduce tags** | |
| ssrA_WT_FP | GC**AGCAAACGACGAAAACTACGCTTTAGCAGCT**TAAGCTTAATTAGCTGAGTCTAGAGGC |
| ssrA_WT_RP | **AGCTGCTAAAGCGTAGTTTTCGTCGTTTGCT**GCTTTGTACAGCTCATCCATGC |
| DAS_FP | C**AGCAAACGACGAAAACTACGCTGATGCATCT**TAAGCTTAATTAGCTGAGTCTAGAGGC |
| DAS_RP | **AGATGCATCAGCGTAGTTTTCGTCGTTTGCT**GCTTTGTACAGCTCATCCATGC |
| DASplus4_FP | **GCAGCAAACGACGAAAACTACTCTGAAAATTATGCTGATGCATCT**TAAGCTTAATTAGCTGAGTCTAGAGGC |
| DASplus4_RP | **AGATGCATCAGCATAATTTTCAGAGTAGTTTTCGTCGTTTGCT**GCTTTGTACAGCTCATCCATGC |
| **qPCR primers** | |
| qPCR_FP | GCATTTATCAGGGTTATTGTCTCAT |
| qPCR_RP | GGGAAATGTGCGCGGAAC |
